# First documentation of *Buprestis haemorrhoidalis* (Coleoptera: Buprestidae) occurrence and damage in Korean wooden cultural heritage: population genetic structure and conservation implications

**DOI:** 10.64898/2026.11.17.767259

**Authors:** Donguk Kim, KyoungSu Choi, Kwang Shik Choi

## Abstract

*Buprestis haemorrhoidalis* Herbst, 1780 (Coleoptera: Buprestidae) is a Palaearctic wood-boring beetle associated with conifers and capable of developing in structural timber. Although this species is known to occur in Korea, its occurrence in wooden cultural heritage structures and its wood-boring activity have not previously been documented or investigated. We surveyed 58 sites in various regions of South Korea, including wooden cultural heritage sites, some designated as UNESCO World Heritage properties, and natural and timber-associated environments. We also examined mitochondrial genetic variation using maximum-likelihood phylogenetic analysis and haplotype networks based on cytochrome *c* oxidase subunit I (*COI*) sequences from Korean specimens and reference sequences from Japan, the Russian Far East, and Europe. Adults were observed in natural and timber-associated environments, while adults and characteristic emergence holes were recorded at wooden cultural heritage sites, including the direct observation of an adult emerging from a wooden column. Analysis of 82 *COI* sequences identified 49 haplotypes and three mitochondrial subgroups (A–C). Subgroups A and B comprised predominantly Korean specimens and co-occurred at three sampling sites. Several haplotypes were shared among geographically separated Korean localities. This study provides the first documentation of *B. haemorrhoidalis* occurrence and associated evidence of wood-boring damage in Korean wooden cultural heritage, together with baseline information on its occurrence in human-made wooden environments and mitochondrial genetic variation. Shared *COI* haplotypes and historical timber-supply patterns raise the possibility of timber-mediated dispersal. These findings provide a foundation for monitoring this species and informing the conservation of wooden cultural heritage in Korea.

## Introduction

*Buprestis haemorrhoidalis* Herbst, 1780 (Coleoptera: Buprestidae) is widely distributed in the Palaearctic region, where it is associated with coniferous trees and recognized as a wood-boring pest (Ekbom 1959, Bílý 2002, Taniwaki 2006, Bellamy 2008, Kubáň 2016). Its larvae develop in dead or dying conifers, including *Picea* and *Abies*, with a preference for *Pinus*, and also utilize stumps, fallen wood, and processed timber. Larval development typically takes two to three years (Bílý 1982, 2002).

In Europe, *B. haemorrhoidalis* has also been documented as a pest of wooden structures. Ekbom (1959) documented adults emerging from building timber and larvae inhabiting wooden structural components in Finland. Dominik (1980) reported damage to wooden buildings, posts, railway sleepers, and bridges in Scandinavia, as well as to wooden structures in Poland. In Japan, Taniwaki (2006) investigated the distribution of emergence holes of this species in *Pinus densiflora* logs. These records indicate that *B. haemorrhoidalis* can infest both natural coniferous substrates and timber used in wooden structures.

Wooden architectural heritage in Korea is susceptible to damage by various biological agents. In particular, termite damage has received considerable attention and has been extensively studied in conservation research (Han and Lee 1990, Kim et al. 2021, Kim and Chung 2022, Kim and Kim 2023, 2024, Im and Han 2024). In contrast, studies on wood-boring beetles have been relatively limited, although several studies have documented coleopteran species associated with historical wooden buildings and damage to wooden artifacts (Han and Lee 1990, Jeong et al. 2003, Cha et al. 2024). *Buprestis haemorrhoidalis* has been recorded from the Korean Peninsula (Bellamy 2008, Kubáň 2016).However, buprestid beetles have not previously been investigated as pests of Korean wooden architectural heritage, and the occurrence and associated wood-boring activity of *B. haemorrhoidalis* remain undocumented in this context.

Despite its broad Palaearctic distribution, the intraspecific mitochondrial *COI* variation and haplotype relationships of *B. haemorrhoidalis* also remain unexplored. To address these knowledge gaps, we conducted field surveys at 58 sites in various regions of South Korea, including wooden cultural heritage sites and natural and timber-associated environments. We investigated the occurrence of *B. haemorrhoidalis* through observations of adults and emergence holes, with particular emphasis on its association with wooden cultural heritage. We also analyzed mitochondrial cytochrome *c* oxidase subunit I (*COI*) sequences using maximum-likelihood phylogenetic inference and haplotype network analysis, incorporating publicly available sequences from Japan, the Russian Far East, and Europe. By integrating field observations with molecular analyses, this study aims to provide the first documentation of *B. haemorrhoidalis* occurrence and associated damage in Korean wooden cultural heritage, establish baseline information on its mitochondrial genetic variation, and provide a foundation for future monitoring and pest management strategies.

## Materials and methods

### Study sites and field surveys

Field surveys were conducted at 58 sites in various regions of South Korea between July 2017 and August 2026 (Fig. 1). Survey sites included wooden cultural heritage sites, some associated with UNESCO World Heritage properties (Fig. 2), and other wooden buildings, such as accommodation facilities and restaurants. Natural and timber-associated environments, as well as outdoor wooden structures, were also surveyed (Fig. 3). Detailed survey records, including localities, dates, site characteristics, numbers of adults observed, and evidence of emergence holes, are provided in Supplementary Tables S1 and S2. The site numbers in Fig. 1 correspond to those listed in Supplementary Tables S1 and S2. Field photographs of surveyed structures, adults, and emergence holes are provided in Supplementary Fig. S1.

**Fig. 1.**
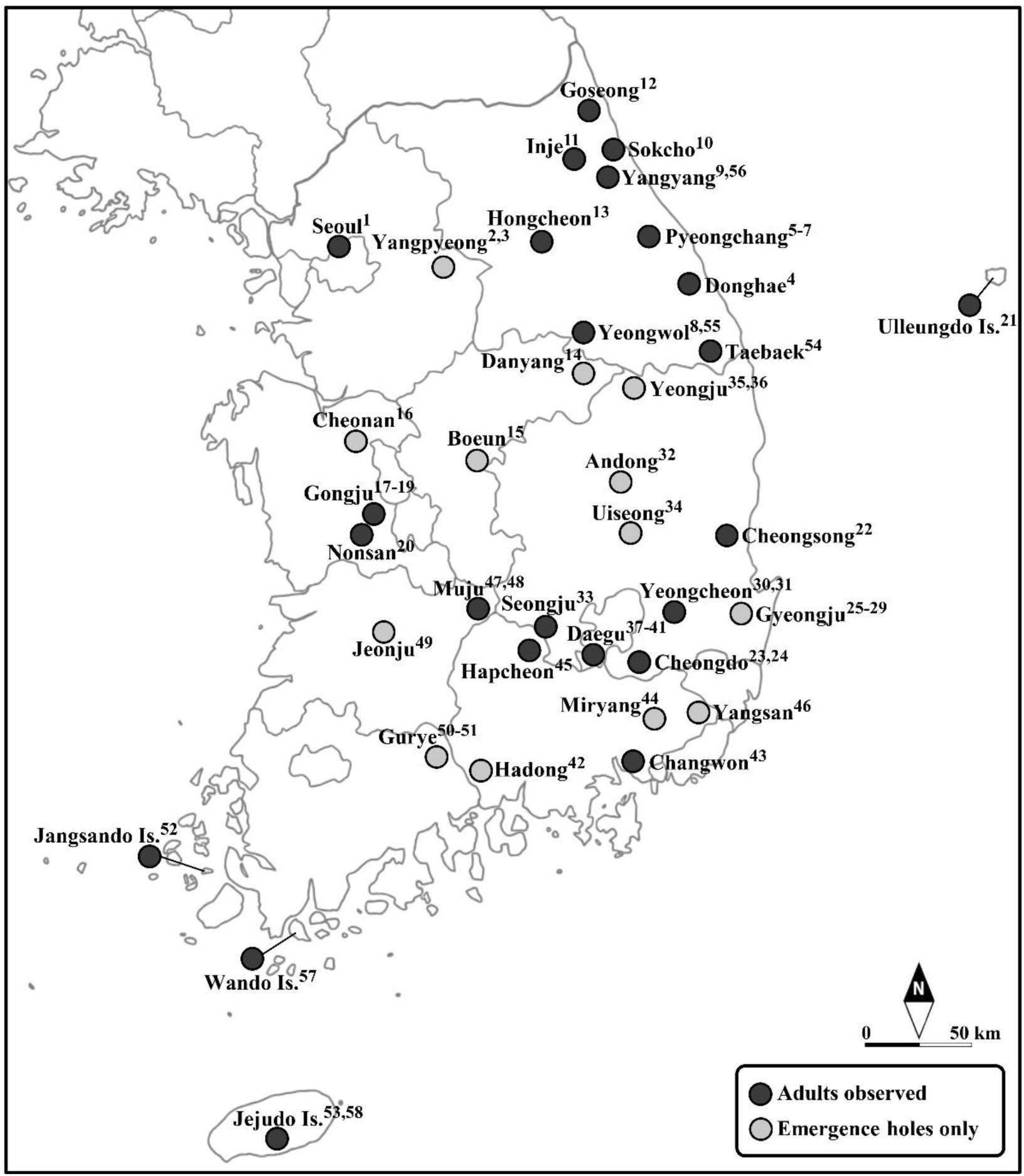
Surveyed localities of *Buprestis haemorrhoidalis* in Korea. Black circles indicate sites where adults were observed, including both live and dead individuals, whereas gray circles indicate sites where only emergence holes were observed. Numbers correspond to the survey sites listed in Supplementary Tables S1 and S2.

**Fig. 2.**
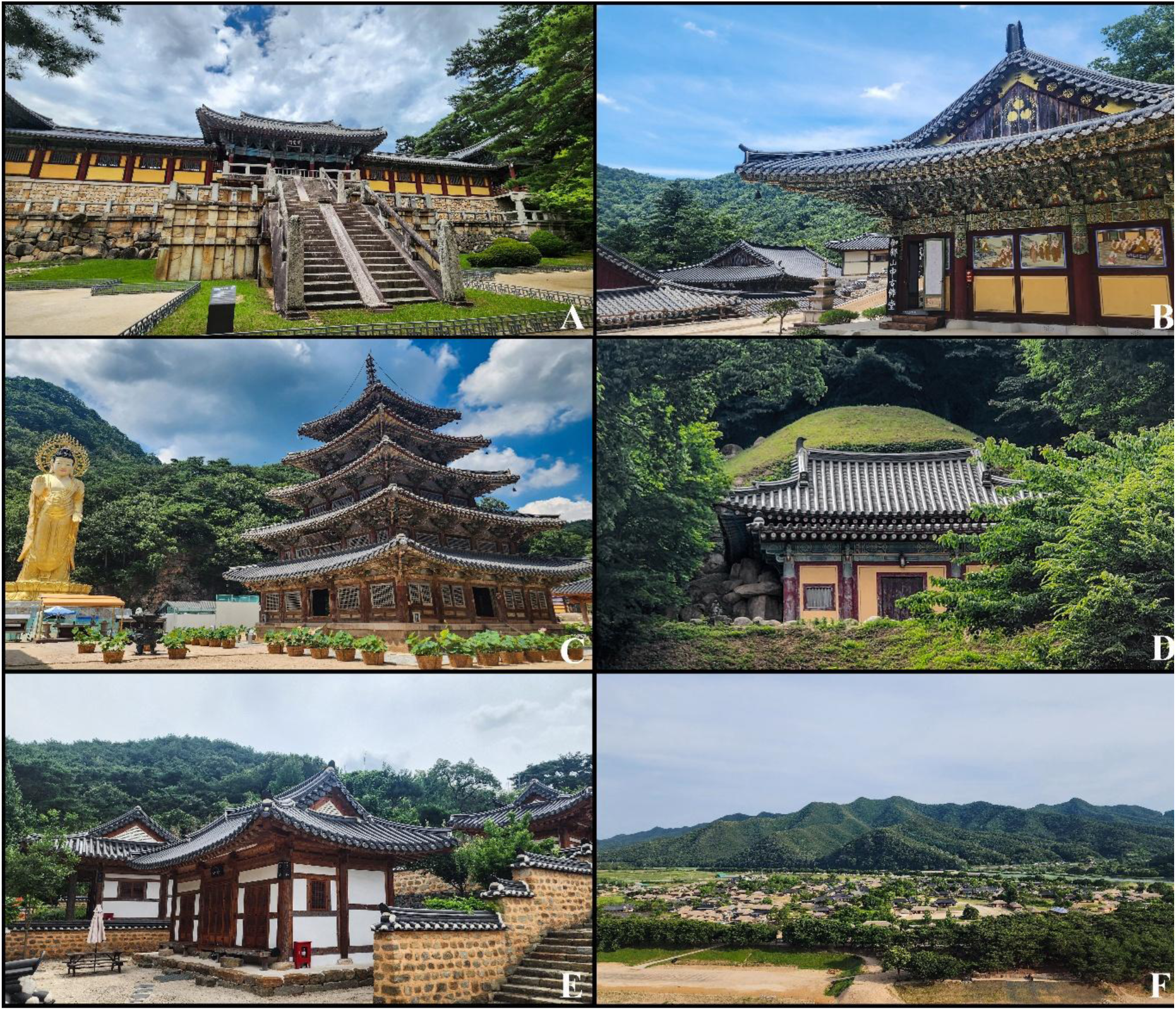
Representative UNESCO World Heritage sites surveyed for *Buprestis haemorrhoidalis* in Korea. (A) Bulguksa Temple, Gyeongju. (B) Haeinsa Temple, Hapcheon. (C) Beopjusa Temple, Boeun. (D) Seokguram Grotto, Gyeongju. (E) Dodongseowon Confucian Academy, Daegu. (F) Hahoe Village, Andong.

**Fig. 3.**
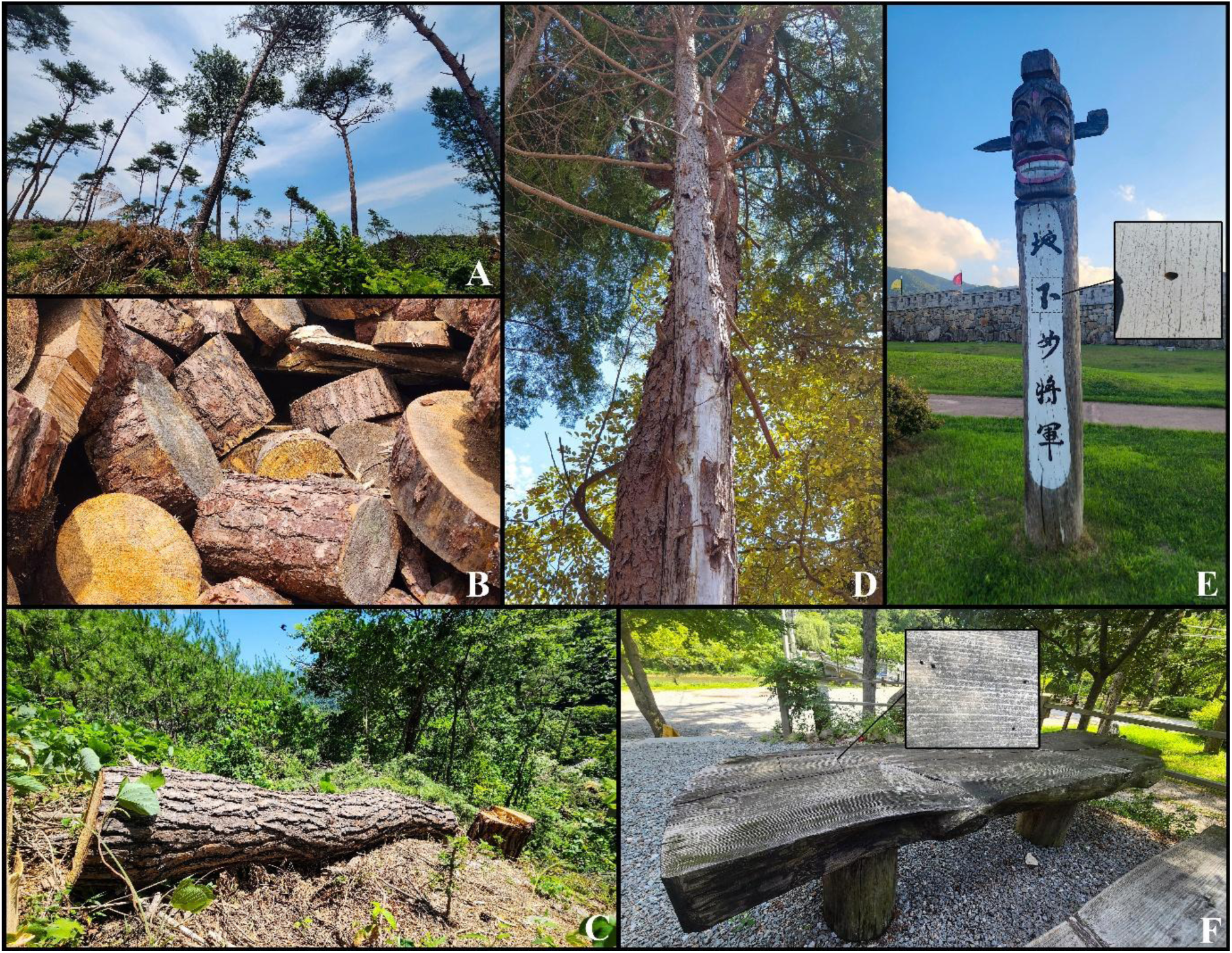
Representative natural and artificial wooden substrates and habitats where *Buprestis haemorrhoidalis* was observed or collected. (A) Logging site. (B) Stacked pine logs. (C) Felled pine trees. (D) Standing dead cypress tree where an adult was collected. (E) Wooden jangseung. (F) Outdoor wooden table.

### Visual inspection and species identification

Adults and emergence holes were surveyed through direct visual inspection of wooden structures and woody substrates. Adult specimens were identified as *Buprestis haemorrhoidalis* based on morphological characters (Bílý 1982, Ohmomo and Fukutomi 2013). Identification was performed at the species level without subspecific assignment. Emergence holes were attributed to *B. haemorrhoidalis* based on their characteristic transversely elliptical shape, with reference to the morphology and dimensions described by Taniwaki (2006). This attribution was further supported by the direct observation of an adult emerging from a wooden column at Seongjusa Temple in Changwon, as well as by dead adults found trapped in or within emergence holes. The number of adults recorded at each survey site included both living and dead individuals, and the presence of emergence holes was also documented. Voucher specimens are deposited in the Donguk Kim Collection (DKCK; Daegu, Republic of Korea), with specimen information provided in Supplementary Table S3.

### DNA extraction, PCR amplification and sequencing

Adult specimens were stored at −70 °C until DNA extraction. Total genomic DNA was extracted from thoracic muscle or membranous tissue surrounding the genitalia of male and female specimens using the DNeasy Blood & Tissue Kit (QIAGEN, Hilden, Germany), following the manufacturer’s protocol. A fragment of the mitochondrial cytochrome *c* oxidase subunit I (*COI*) gene was amplified by polymerase chain reaction (PCR) using one of two primer pairs: LepF1 (5′-ATTCAACCAATCATAAAGATATTGG-3′) and LepR1 (5′-TAAACTTCTGGATGTCCAAAAAATCA-3′) (Hebert et al. 2004), or LCO1490 (5′-GGTCAACAAATCATAAAGATATTGG-3′) and HCO2198 (5′-TAAACTTCAGGGTGACCAAAAAATCA-3′) (Folmer et al. 1994). One primer pair was used for each specimen, and the same PCR conditions were applied to both primer pairs. PCR was performed in a total volume of 25 μL, consisting of 24 μL of reaction mixture (1× PCR buffer, 0.2 mM dNTPs, 0.4 μM of each primer, and 0.5 U of Taq DNA polymerase [TaKaRa, Shiga, Japan]) and 1 μL of genomic DNA. The thermal cycling conditions consisted of an initial denaturation at 94 °C for 2 min, followed by 35 cycles of denaturation at 94 °C for 1 min, annealing at 48 °C for 1 min, and extension at 72 °C for 1 min, with a final extension at 72 °C for 5 min. PCR products were examined by electrophoresis on a 1.5% agarose gel, purified, and sequenced by Macrogen (Daejeon, Republic of Korea). Raw sequences were assembled and edited using BioEdit (Hall 1999). GenBank accession numbers for the *COI* sequences generated in this study are provided in Supplementary Table S3.

### Sequence alignment, phylogenetic analysis, and haplotype network analysis

A total of 76 *COI* sequences were newly generated from Korean specimens of *Buprestis haemorrhoidalis* and analyzed together with six publicly available reference sequences from the Russian Far East (n = 1), Japan (n = 1), and Europe (n = 4; Finland, Poland, and Italy), retrieved from NCBI GenBank and the Barcode of Life Data System (BOLD). A *COI* sequence of *B. splendens* was included as an outgroup. Sequences were aligned using ClustalW implemented in MEGA12 (Kumar et al. 2024), and the alignment was trimmed to 658 bp to standardize sequence length. The final dataset comprised 83 sequences, including 82 *B. haemorrhoidalis* sequences and one outgroup sequence. Geographic origins of the sequences are provided in Supplementary Tables S3.

Phylogenetic relationships were inferred using maximum-likelihood (ML) and neighbor-joining (NJ) methods. The ML analysis was performed using IQ-TREE 3.1.3 (Wong et al. 2026). The best-fitting nucleotide substitution model (TN+F+I+R2) was selected using ModelFinder (Kalyaanamoorthy et al. 2017) based on the Bayesian information criterion (BIC). Branch support was assessed using the SH-like approximate likelihood ratio test (SH-aLRT) with 1,000 replicates and ultrafast bootstrap analysis with 1,000 replicates (Hoang et al. 2018). The NJ analysis was performed using MEGA12 under the Kimura two-parameter (K2P) model, with branch support evaluated using 1,000 bootstrap replicates. The resulting ML tree was visualized using FigTree v1.4.4 (Rambaut 2018).

Mean pairwise genetic distances within and between mitochondrial subgroups A–C, identified in the ML phylogeny, were calculated under the K2P model using MEGA12. All 82 *B. haemorrhoidalis COI* sequences were included in the distance analysis, excluding the outgroup. Mean within- and between-subgroup distances were calculated by averaging pairwise distances between sequences within each subgroup and between sequences from different subgroups, respectively. A median-joining (MJ) haplotype network was constructed from the same 82 sequences using PopART (Leigh and Bryant 2015), following the algorithm of Bandelt et al. (1999). The resulting network was visualized in two ways, with haplotypes colored according to mitochondrial subgroup assignments (A–C) or geographic origins. Haplotype circles were colored according to the respective categories, with circle sizes proportional to the number of specimens sharing each haplotype.

## Results

### Occurrence and distribution of *Buprestis haemorrhoidalis*

Field surveys at 58 sites in various regions of South Korea documented the occurrence of *Buprestis haemorrhoidalis* in wooden cultural heritage sites, natural and timber-associated environments, and other wooden structures (Fig. 1, Supplementary Tables S1 and S2). At cultural heritage sites, evidence of occurrence included adults, characteristic emergence holes, or both. The surveyed cultural heritage sites included Buddhist temples, Confucian academies (Seowon), and traditional villages, some associated with UNESCO World Heritage properties (Fig. 2). Other survey sites included historic and commercial wooden buildings, such as accommodation facilities and restaurants. Detailed occurrence records are provided in Supplementary Tables S1 and S2, and corresponding field photographs are presented in Supplementary Fig. S1.

Adults were also recorded in natural and timber-associated environments, including logging areas, stacked and felled logs, and standing dead trees (Fig. 3A–D). Characteristic emergence holes were also observed in outdoor wooden structures, including traditional wooden jangseung and outdoor tables (Fig. 3E, F). These observations indicate that this species occurs in a variety of natural and human-made woody environments in South Korea.

### Emergence holes and associated damage to wooden structures

Emergence holes attributed to *Buprestis haemorrhoidalis* were observed in wooden structures at surveyed cultural heritage sites. The holes were characteristically transversely elliptical and occurred in wooden components showing varying degrees of weathering, including columns of old temple buildings (Fig. 4A, B). This attribution was further supported by dead adults found trapped in emergence holes or within holes with their pronota partially exposed (Fig. 4C, D). However, the timing of formation of most emergence holes could not be determined from their external appearance.

**Fig. 4.**
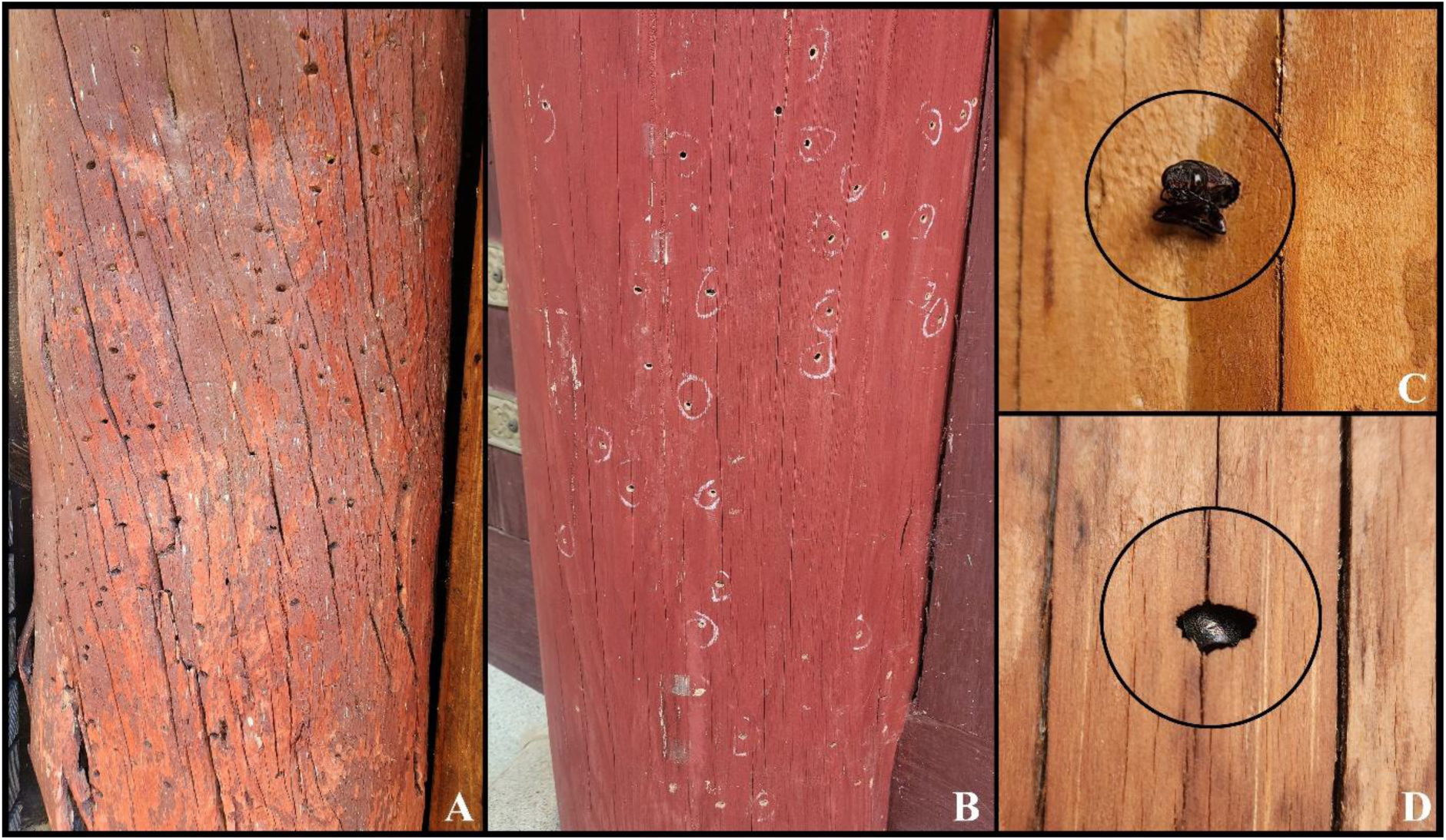
Emergence holes attributed to *Buprestis haemorrhoidalis* and associated adult remains in wooden structures. (A) Emergence holes on a weathered wooden column at Seongjusa (Changwon). (B) Emergence holes marked with white circles on a painted wooden column at Woljeongsa (Pyeongchang). (C) A dead adult trapped in an emergence hole at Sinheungsa (Sokcho). (D) A dead adult found within an emergence hole, with the pronotum partially exposed, at Sinheungsa (Sokcho).

Emergence holes were observed in various architectural components of wooden buildings, including columns (Fig. 5A, B, E, F), floorboards (Fig. 5C), rafters (Fig. 5D), doors (Fig. 5G), and thresholds (Fig. 5H).

**Fig. 5.**
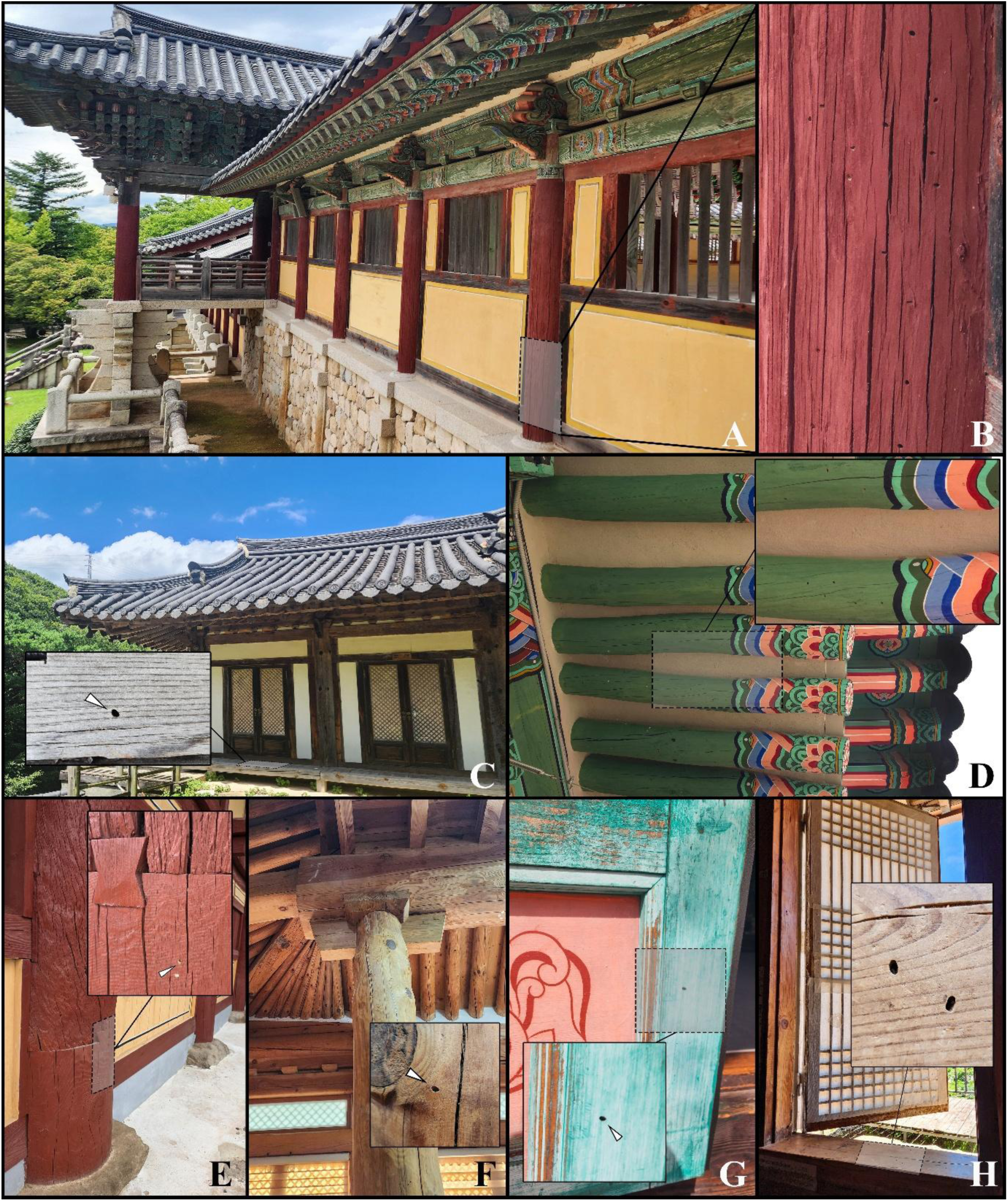
Emergence holes attributed to *Buprestis haemorrhoidalis* in various wooden building components. (A) Wooden columns at Bulguksa (Gyeongju). (B) Close-up view of the emergence holes shown in (A). (C) Wooden floorboards at Seongjusa (Changwon). (D) Wooden rafters at Eunhaesa (Yeongcheon). (E) Repaired wooden column at Eunhaesa (Yeongcheon). (F) Interior wooden column of a commercial building. (G) Wooden doors at Gounsa (Uiseong). (H) Wooden threshold at an accommodation facility. White arrows indicate emergence holes, and enlarged views show their details.

### Adult behavior and secondary occupation of emergence holes

Oviposition behavior of female *Buprestis haemorrhoidalis* was observed in cracks and crevices of wooden structures (Fig. 6A–C). An adult was also observed emerging from a wooden column at Seongjusa Temple in Changwon (Fig. 6D), providing direct evidence of adult emergence from structural timber. Potter wasps (Hymenoptera: Vespidae: Eumeninae) were observed using emergence holes attributed to *B. haemorrhoidalis* as nesting sites, with individuals entering and exiting the holes (Fig. 6E, F). These observations indicate that such holes can subsequently serve as nesting sites for other insects.

**Fig. 6.**
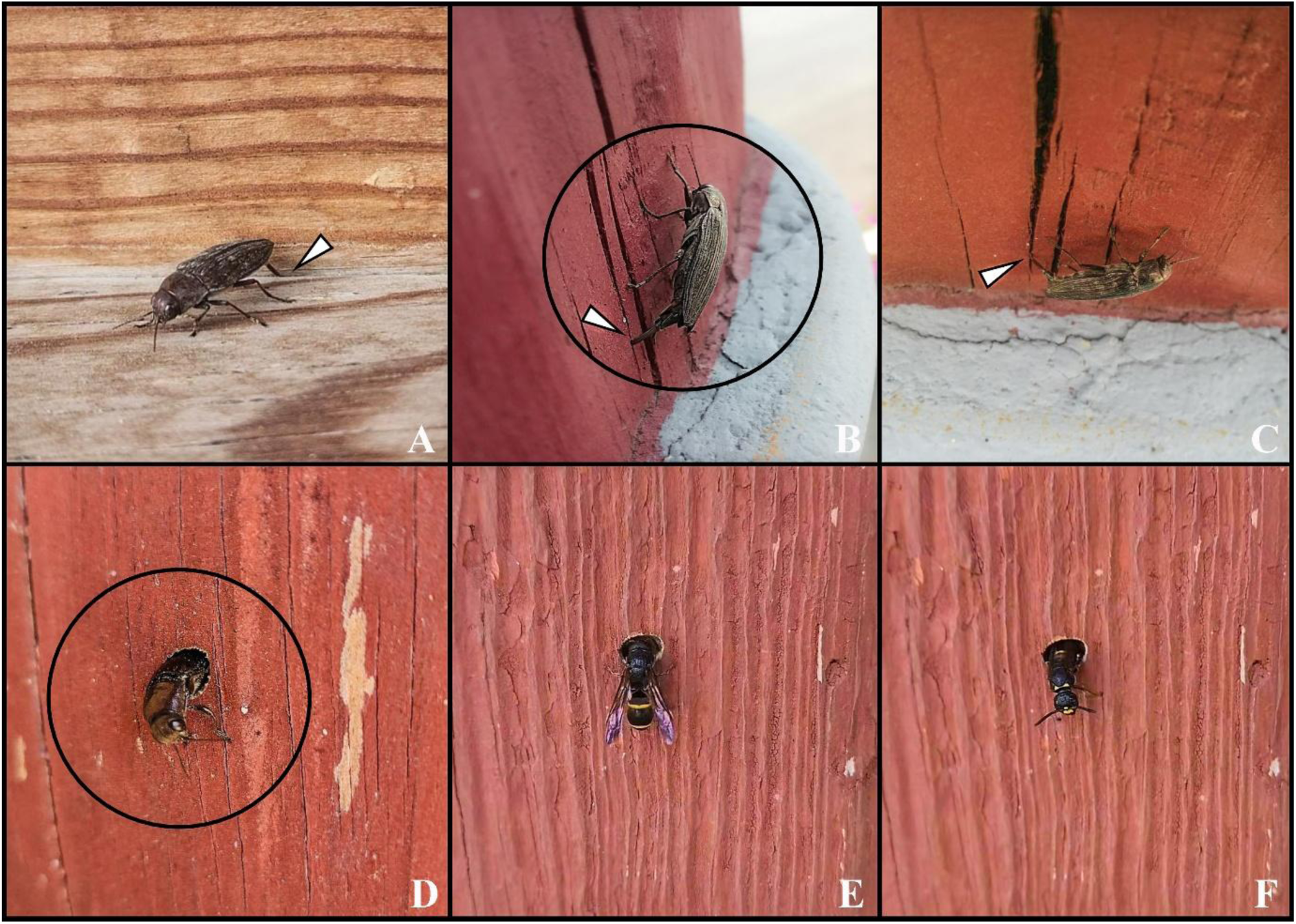
Behavioral observations of *Buprestis haemorrhoidalis* and secondary use of emergence holes by potter wasps (Hymenoptera: Vespidae: Eumeninae). (A–C) Oviposition behavior of female *B. haemorrhoidalis* in cracks and crevices of wooden structures at Yugasa (Daegu) (A) and Samhwasa (Donghae) (B, C). (D) An adult *B. haemorrhoidalis* emerging from a wooden column at Seongjusa (Changwon). (E, F) Potter wasps entering (E) and exiting (F) emergence holes used as nesting sites at Eunhaesa (Yeongcheon).

### Mitochondrial *COI* variation and phylogenetic relationships

Maximum-likelihood (ML) and neighbor-joining (NJ) analyses of the 658-bp *COI* dataset recovered three major mitochondrial subgroups, designated A, B, and C (Fig. 7A). Subgroup A comprised 45 specimens, including 44 from South Korea and one reference sequence from the Russian Far East. Subgroup B comprised 33 specimens, including 32 from South Korea and one reference sequence from Japan. Subgroup C comprised four European reference sequences from Finland, Poland, and Italy (Fig. 7A, B, Supplementary Table S3). The ML and NJ analyses recovered congruent major mitochondrial subgroups.

**Fig. 7.**
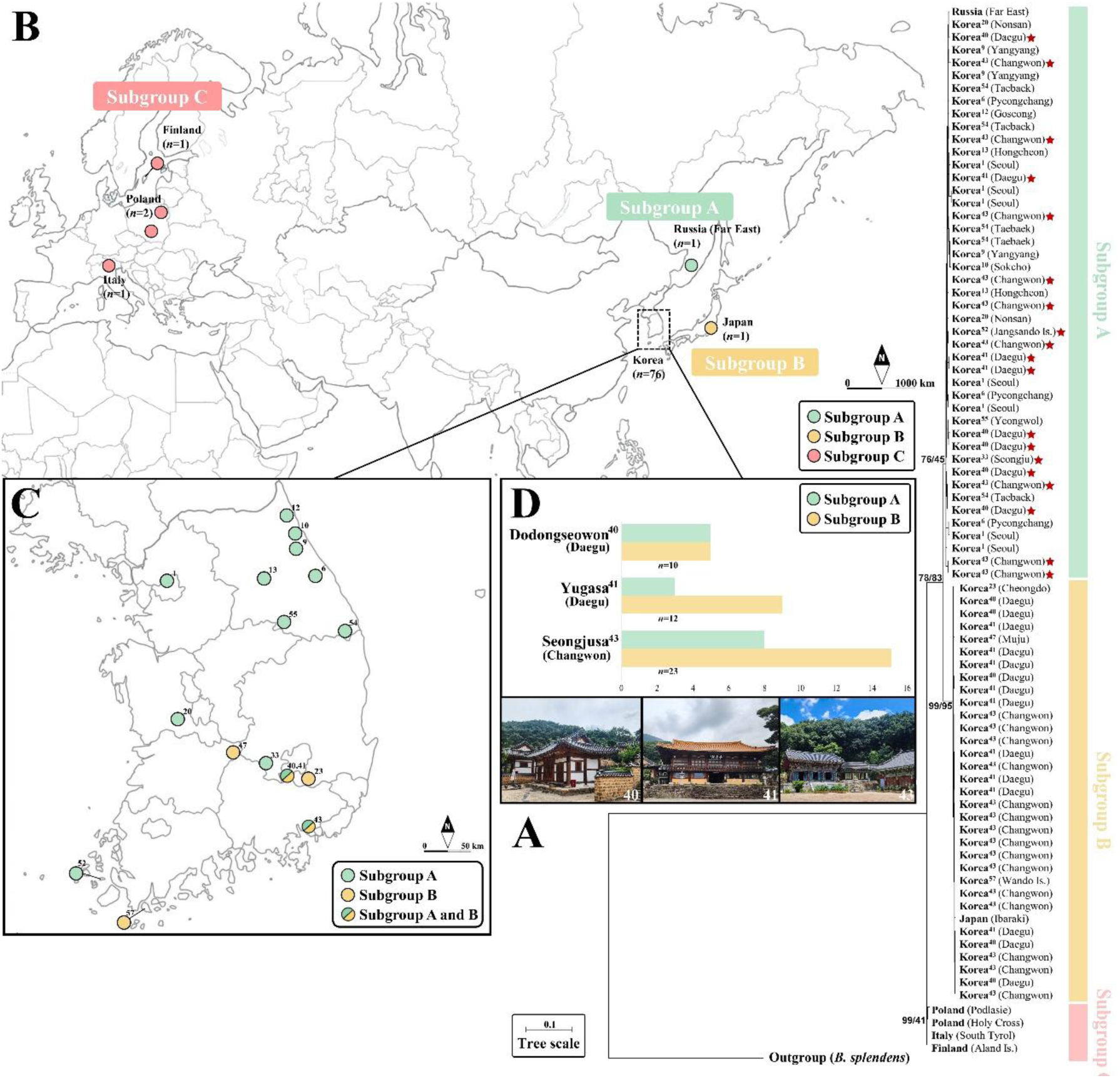
Mitochondrial *COI* phylogenetic relationships and geographic origins of *Buprestis haemorrhoidalis* specimens included in the analyses. (A) Maximum-likelihood (ML) phylogenetic tree showing three mitochondrial subgroups (A–C), with *B. splendens* as the outgroup. Numbers at major nodes indicate neighbor-joining (NJ) bootstrap and ML ultrafast bootstrap support values, respectively (NJ/ML). Red stars indicate specimens collected from southern Korean localities that were assigned to subgroup A. (B) Geographic origins of the *COI* sequences included in the phylogenetic analyses, colored according to mitochondrial subgroup. (C) Sampling localities of South Korean specimens included in the *COI* analyses, colored according to mitochondrial subgroup (A or B). Bicolored symbols indicate sampling sites where specimens belonging to both subgroups were detected. (D) Numbers of specimens assigned to subgroups A and B at three sites where both subgroups occurred: Dodongseowon (Daegu), Yugasa (Daegu), and Seongjusa (Changwon). Photographs below the bar charts show the corresponding survey sites in the same order. Green, yellow, and pink indicate subgroups A, B, and C, respectively. Superscript numbers indicate survey site numbers corresponding to those in Fig. 1 and Supplementary Tables S1 and S2.

Korean specimens assigned to subgroups A and B were collected from overlapping geographic areas (Fig. 7C, Supplementary Table S3). Subgroup A specimens originated from multiple regions of South Korea, whereas subgroup B specimens were collected predominantly from southern localities. Although most specimens from southern localities belonged to subgroup B, 19 specimens from these areas were assigned to subgroup A (Fig. 7A, C). Both subgroups were detected at three surveyed sites: Dodongseowon and Yugasa in Daegu, and Seongjusa in Changwon. At Dodongseowon, five specimens belonged to each subgroup. At Yugasa, three specimens belonged to subgroup A and nine to subgroup B, whereas at Seongjusa, eight belonged to subgroup A and 15 to subgroup B (Fig. 7D). These results confirm the co-occurrence of the two mitochondrial subgroups at three surveyed sites.

Mean pairwise Kimura two-parameter (K2P) genetic distances between subgroups were 2.98% for A–B, 4.18% for A–C, and 4.75% for B–C (Supplementary Table S4). Mean within-subgroup distances were 0.82%, 0.24%, and 0.41% for subgroups A, B, and C, respectively. The lowest mean between-subgroup distance was observed between subgroups A and B.

### Haplotype diversity and network relationships

Analysis of 82 *COI* sequences of *Buprestis haemorrhoidalis* identified 49 haplotypes, including 36 in subgroup A, 10 in subgroup B, and three in subgroup C. When colored according to the mitochondrial subgroups identified in the maximum-likelihood (ML) phylogeny, the median-joining network revealed three distinct haplotype assemblages (Fig. 8A). Subgroup A contained more haplotypes and exhibited a more extensively branched network structure than subgroup B, whereas subgroup C comprised three haplotypes represented by European reference sequences.

**Fig. 8.**
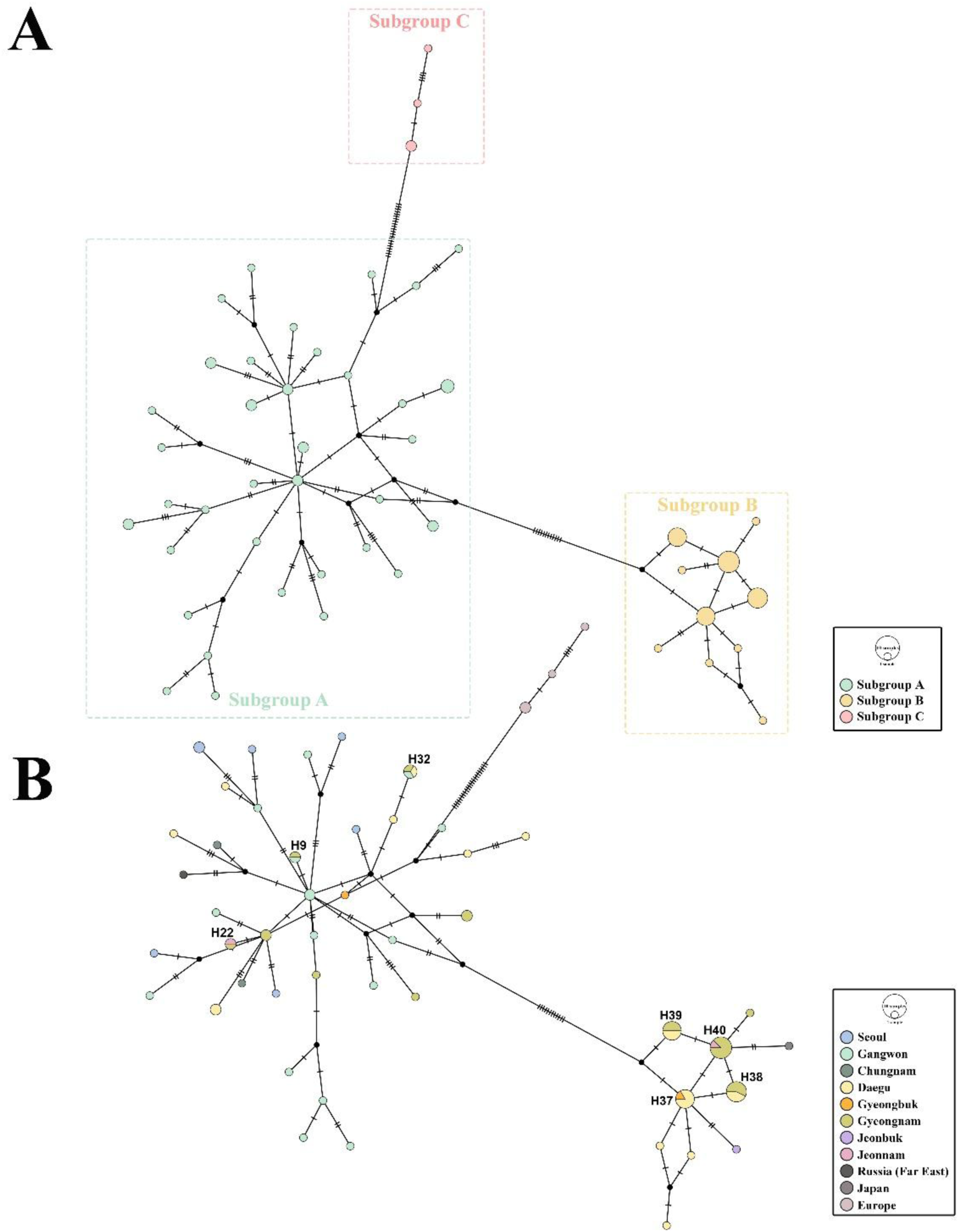
Median-joining haplotype network based on 82 mitochondrial *COI* sequences of *Buprestis haemorrhoidalis*, visualized using two different color schemes. (A) Network colored according to mitochondrial subgroups A–C identified in the maximum-likelihood phylogeny. Dashed boxes indicate the three mitochondrial subgroups. (B) The same network colored according to geographic origin. Seven haplotypes shared among geographic regions (H9, H22, H32, and H37–H40) are labeled. Multicolored circles represent haplotypes shared by specimens from different geographic regions. Circle sizes indicate haplotype frequencies, hatch marks indicate mutational steps, and black nodes represent inferred median vectors.

Visualization of the same network according to geographic origin revealed seven haplotypes shared among different regions of South Korea (Fig. 8B). Three belonged to subgroup A: H9 was shared between Gangwon and Gyeongnam, H22 between Jeonnam and Gyeongnam, and H32 among Gangwon, Daegu, and Gyeongnam. The remaining four belonged to subgroup B: H37 was shared between Daegu and Gyeongbuk, H38 and H39 were each shared between Daegu and Gyeongnam, and H40 between Jeonnam and Gyeongnam. These results indicate that identical *COI* haplotypes occurred across geographically separated sampling regions within both subgroups A and B.

## Discussion

### Occurrence and damage associated with wooden cultural heritage

The present study provides systematic field evidence of the occurrence of *Buprestis haemorrhoidalis* in association with wooden cultural heritage and other wooden structures in Korea. Characteristic emergence holes were observed in various architectural components, including columns, floorboards, rafters, doors, and thresholds (Fig. 5). Adult specimens were also found in natural and timber-associated environments, including logging sites, woodpiles, and standing dead trees (Fig. 3). These observations indicate that the species occurs in a range of woody environments, including both natural substrates and human-made structures.

The emergence holes observed in wooden buildings are consistent with the wood-boring biology of *B. haemorrhoidalis*. Ekbom (1959) documented the emergence of this species from structural timber in Finnish buildings, including wooden window components, while Taniwaki (2006) described its characteristic transversely elliptical emergence holes in *Pinus densiflora* logs in Japan. In the present study, the attribution of emergence holes to *B. haemorrhoidalis* was further supported by direct observation of an adult emerging from a wooden column of a Buddhist temple building (Fig. 6D), as well as by dead adults found in or trapped within emergence holes (Fig. 4C, D). Nevertheless, the presence of emergence holes does not necessarily indicate an active infestation, as the precise timing of their formation could not be determined for most holes observed in this study.

Such holes are consistent with previous adult emergence of *Buprestis haemorrhoidalis* and may indicate the presence of larval galleries within the wood. Their occurrence in various architectural components therefore warrants attention in the conservation of wooden cultural heritage. However, the present study did not assess the extent of internal wood damage or its effects on structural integrity. Accordingly, the observed emergence holes should be regarded as evidence of past wood-boring activity attributed to *B. haemorrhoidalis*, rather than as indicators of current infestation or the severity of structural damage. Furthermore, potter wasps (Hymenoptera: Vespidae: Eumeninae) were observed using emergence holes attributed to *B. haemorrhoidalis* as nesting sites (Fig. 6E, F), demonstrating that these holes can subsequently be used by other insects. This secondary use further emphasizes the need to distinguish the original wood-boring activity from subsequent occupation of existing emergence holes.

### Mitochondrial diversity and geographic overlap of subgroups

Phylogenetic analyses of mitochondrial *COI* sequences identified three major subgroups of *Buprestis haemorrhoidalis* (Fig. 7A). Subgroups A and B comprised predominantly Korean specimens, together with one reference sequence from the Russian Far East and one from Japan, respectively, whereas subgroup C comprised four European reference sequences. The mean K2P genetic distance between subgroups A and B was 2.98%, lower than those between A and C (4.18%) and between B and C (4.75%) (Supplementary Table S4). These results indicate mitochondrial differentiation among the analyzed specimens, although the limited geographic representation of the reference sequences precludes a comprehensive assessment of genetic variation across the species’ broader geographic range.

Subgroup A comprised 36 haplotypes among 45 specimens, compared with 10 haplotypes among 33 specimens in subgroup B (Fig. 8). The greater haplotype richness and higher mean within-subgroup K2P distance observed in subgroup A indicate greater mitochondrial variation among the sampled specimens of this subgroup. However, differences in sample size and geographic representation should be considered when comparing the diversity of the two subgroups. Furthermore, the four European reference sequences assigned to subgroup C are insufficient to characterize mitochondrial diversity across European populations.

Although subgroup B was found predominantly in southern Korea, specimens belonging to both subgroups A and B were found in some of the same areas. Nineteen specimens collected from southern localities belonged to subgroup A, and both subgroups were detected at three surveyed sites: Dodongseowon, Yugasa, and Seongjusa. These observations demonstrate that the two mitochondrial subgroups co-occurred at several surveyed sites. In addition, the haplotype network revealed seven haplotypes shared among different geographic regions within South Korea, including three in subgroup A and four in subgroup B. Notably, H32 was shared among specimens from Gangwon, Daegu, and Gyeongnam, while other shared haplotypes were detected across different regions within both subgroups. These findings show that some *COI* haplotypes occurred across geographically separated sampling regions within the same mitochondrial subgroup. Nevertheless, neither the local co-occurrence of mitochondrial subgroups nor the sharing of *COI* haplotypes among regions provides direct evidence of contemporary gene flow or interbreeding (Ballard and Whitlock 2004).

### Potential anthropogenic dispersal through timber transportation

The occurrence of identical *COI* haplotypes in geographically separated Korean localities raises the possibility that human-mediated timber movement has contributed to the dispersal of *Buprestis haemorrhoidalis*. Such a process is biologically plausible because the immature stages develop within wood and may remain concealed during timber handling and transportation. Ekbom (1959) documented the emergence of *B. haemorrhoidalis* from structural timber in buildings in Finland, while Taniwaki (2006) reported emergence holes of this species in *Pinus densiflora* logs in Japan. These observations indicate that the species can develop in wood that may subsequently be used or transported by humans, providing a biological basis for considering timber transportation as a potential dispersal pathway.

Historical and modern timber procurement practices in Korea provide further context for this hypothesis. Pine has long been an important construction material in Korean wooden architecture, with Gangwon and Gyeongbuk serving as major timber-supply regions for historical buildings and wooden cultural heritage (Kim 2005, Park and Lee 2007). According to historical production data summarized by Kim (2005), the Gangwon–northern Gyeongbuk east-coast region accounted for 73.8% of the reported pine timber production associated with wooden cultural heritage (Fig. 9A). A newspaper profile also reported that a traditional carpenter involved in construction and repair work at Seongjusa Temple in Changwon operated a timber workshop in Hoengseong, Gangwon, where locally harvested pine was processed (Yoon 2018). Although this account illustrates connections between timber-related activities in different regions, it does not establish that timber from Gangwon was transported to Seongjusa. Nevertheless, these historical and modern practices provide a relevant context for considering interregional timber transportation as a potential pathway for the dispersal of wood-boring insects.

**Fig. 9.**
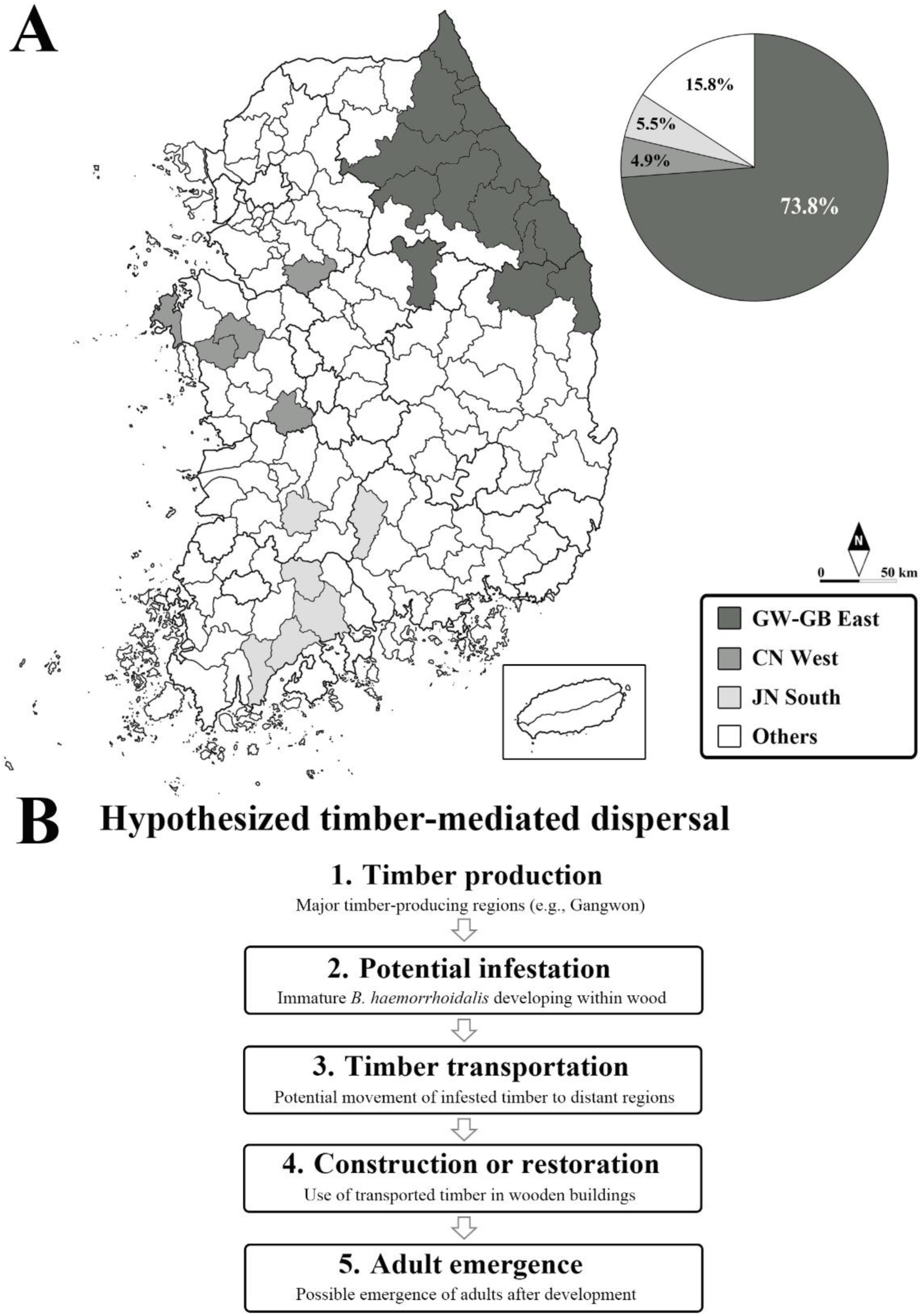
Historical pine timber production and a hypothesized pathway of timber-mediated dispersal of *Buprestis haemorrhoidalis* in Korea. (A) Major historical pine timber production regions associated with wooden cultural heritage, based on data reported by Kim (2005). The pie chart shows the proportions attributed to the Gangwon–northern Gyeongbuk east-coast region (73.8%), Chungnam west-coast region (4.9%), Jeonnam south-coast region (5.5%), and other regions (15.8%). (B) Conceptual diagram illustrating a possible dispersal pathway involving the presence of immature *B. haemorrhoidalis* in timber, transportation to other regions, use in wooden buildings, and subsequent adult emergence. The proposed pathway is hypothetical and was not directly demonstrated in this study.

Within this context, the occurrence of *COI* haplotypes shared between Gangwon and geographically distant Korean localities is consistent with the possibility of timber-mediated dispersal. For example, H32 was shared among specimens from Gangwon, Daegu, and Gyeongnam, while H9 was shared between Gangwon and Gyeongnam. Subgroup A was represented in both northern and southern parts of South Korea, whereas subgroup B was detected predominantly in southern localities and was not found among the specimens sampled from Gangwon, Seoul, or Chungnam. This geographic pattern is noteworthy given the historical concentration of timber production in Gangwon and northern Gyeongbuk. Timber transportation may have facilitated the spread of existing mitochondrial haplotypes to geographically distant regions. Under this hypothetical scenario, immature *B. haemorrhoidalis* developing within harvested timber could be transported to other regions, incorporated into wooden structures, and subsequently emerge as adults after completing development (Fig. 9B). Although timber transportation would not itself generate new mitochondrial haplotypes, repeated introductions of individuals carrying different pre-existing haplotypes could increase haplotype richness at recipient localities. This mechanism may help explain the broad geographic occurrence of subgroup A, but the present mitochondrial data cannot distinguish human-mediated dispersal from natural dispersal.

In addition, other possible explanations should be considered. Shared *COI* haplotypes may reflect historical population connectivity, retention of ancestral mitochondrial variation, or natural dispersal among localities (Knowles 2009). Because no specimens from North Korea were available and only one reference sequence from the Russian Far East was included, the distribution of subgroup B in the northern Korean Peninsula and adjacent regions remains poorly resolved. More broadly, *COI* data alone are insufficient to reliably determine the direction, timing, or mechanism of dispersal (Ballard and Whitlock 2004, Toews and Brelsford 2012).

Inquiries to site administrators and construction contractors regarding the timber used at Dodongseowon, Yugasa, and Seongjusa did not yield verifiable information on its provenance, either because the sources were unknown or no response was received. Consequently, the origin of the timber used at these sites could not be established, and timber-mediated dispersal remains a plausible hypothesis rather than a demonstrated explanation for the observed genetic patterns. Further investigation combining additional genetic markers with timber provenance records and direct evidence of infestation in transported wood would be necessary to evaluate this hypothesis.

### Implications for heritage conservation and future research

The findings of this study have practical implications for the monitoring and conservation of wooden cultural heritage in South Korea. The occurrence of characteristic emergence holes in various architectural components, together with direct observations of adult emergence and oviposition, indicates that wooden structures can provide suitable substrates for the development and reproduction of *Buprestis haemorrhoidalis*. However, existing emergence holes do not necessarily indicate ongoing infestation or reveal the extent of internal wood damage. Conservation assessments should therefore distinguish evidence of past wood-boring activity from current infestation by monitoring newly formed emergence holes and adult emergence, while also considering wood condition and previous repair history.

The potential role of timber transportation in the dispersal of *B. haemorrhoidalis* also highlights the importance of documenting the origin and handling of wood used in construction and restoration. Records of timber provenance, storage conditions, and replacement history could help identify potential pathways for the introduction of wood-boring insects into wooden structures. Such information would also provide valuable context for interpreting the geographic patterns of genetic variation observed in future studies.

The subspecific classification of *B. haemorrhoidalis* in East Asia remains unresolved (Hołyński 2018). Therefore, mitochondrial subgroups A and B identified in this study should not be directly equated with previously described subspecies without further integrative taxonomic investigation. Future research should incorporate more geographically representative sampling and additional genetic markers, particularly nuclear or genome-wide data, to clarify the relationships between mitochondrial subgroups A and B and assess population connectivity. Direct examination of transported and installed timber for immature stages, together with timber provenance records, would be valuable for evaluating the possibility of anthropogenic dispersal. Long-term monitoring of selected wooden buildings could also help distinguish recent emergence activity from historical infestations and determine whether successive generations develop within installed structural timber.

Overall, this study documents the occurrence of *B. haemorrhoidalis* in association with wooden cultural heritage in South Korea and reveals mitochondrial differentiation and haplotype diversity, including the local co-occurrence of two major subgroups and shared *COI* haplotypes across geographically separated regions. These findings provide a basis for further investigation of the species’ dispersal and population structure and support the development of evidence-based monitoring strategies for its occurrence in wooden cultural heritage.

## Acknowledgements

The first author is especially grateful to Dooyoung Kim (University of California, Berkeley, USA) for his substantial contributions to field surveys and for providing valuable specimens. The first author also thanks Dr. Dongmin Kim and Dr. Young-Kun Kim (Kyungpook National University, Daegu, Republic of Korea), and Junhyeong Choi (Seoul National University, Seoul, Republic of Korea) for sharing field-site information and providing specimens, and Boyun Kim and Sohyun Lee (Kyungpook National University, Daegu, Republic of Korea) for providing photographs of emergence holes. We thank the Korea National Park Service (KNPS) for granting permission to collect insects within national parks. ChatGPT (OpenAI) was used to assist with English-language editing and manuscript formatting. All AI-assisted edits were reviewed and verified by the authors.

## Data availability

The *COI* sequences generated in this study have been deposited in GenBank, and their accession numbers are provided in Supplementary Table S3. Survey records are provided in Supplementary Tables S1 and S2. Specimen information, haplotype and mitochondrial subgroup assignments, geographic origins of the analyzed sequences, and mean K2P genetic distances are provided in Supplementary Tables S3 and S4.

## Supplementary Material

Supplementary material is available with this preprint.

