## Supplementary Figure S1 for "First documentation of *Buprestis haemorrhoidalis* (Coleoptera: Buprestidae) occurrence and damage in Korean wooden cultural heritage: population genetic structure and conservation implications"

**Supplementary Fig. S1.** Field photographs of wooden structures, emergence holes, and oviposition behavior of *Buprestis haemorrhoidalis*.

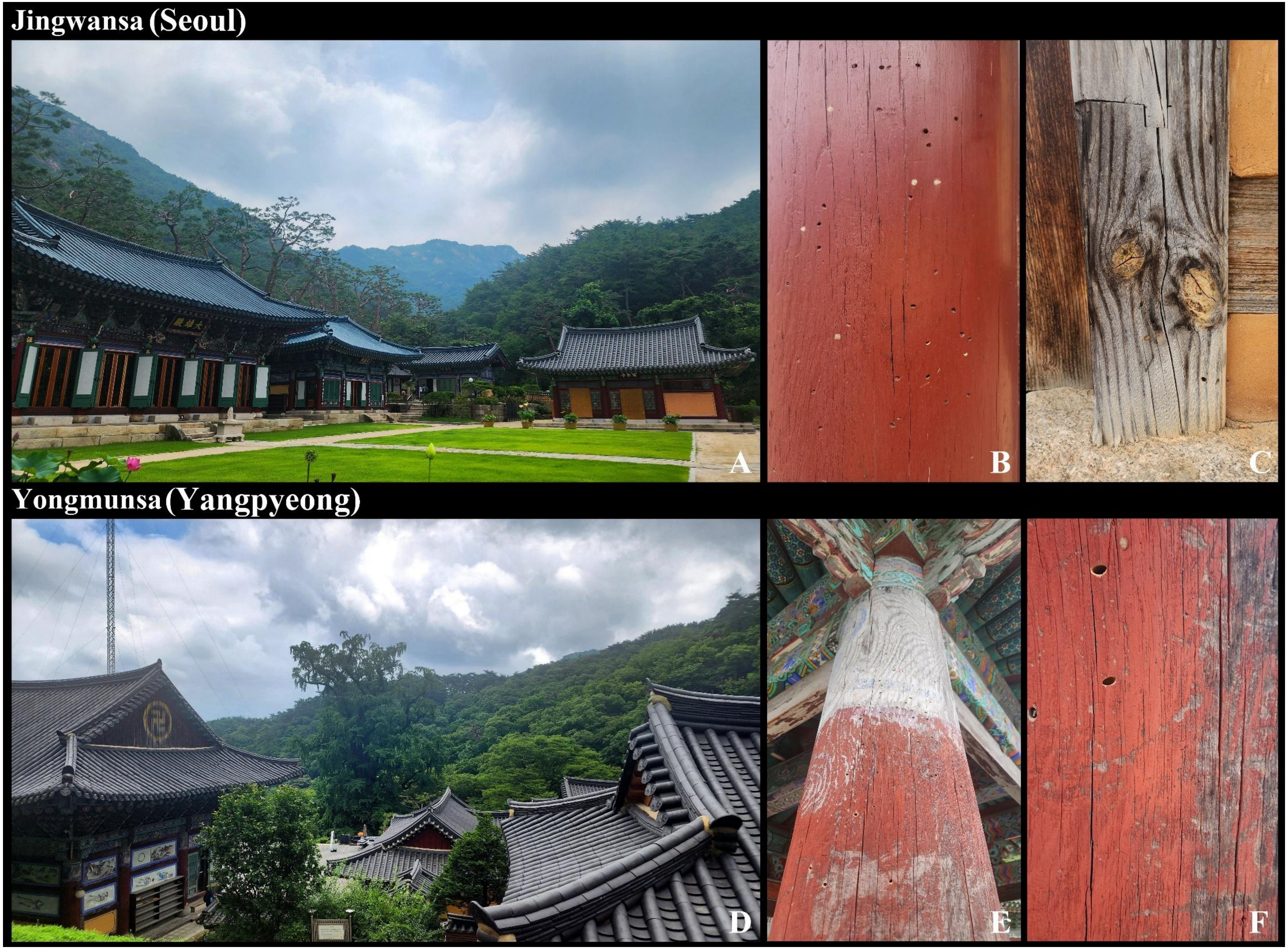

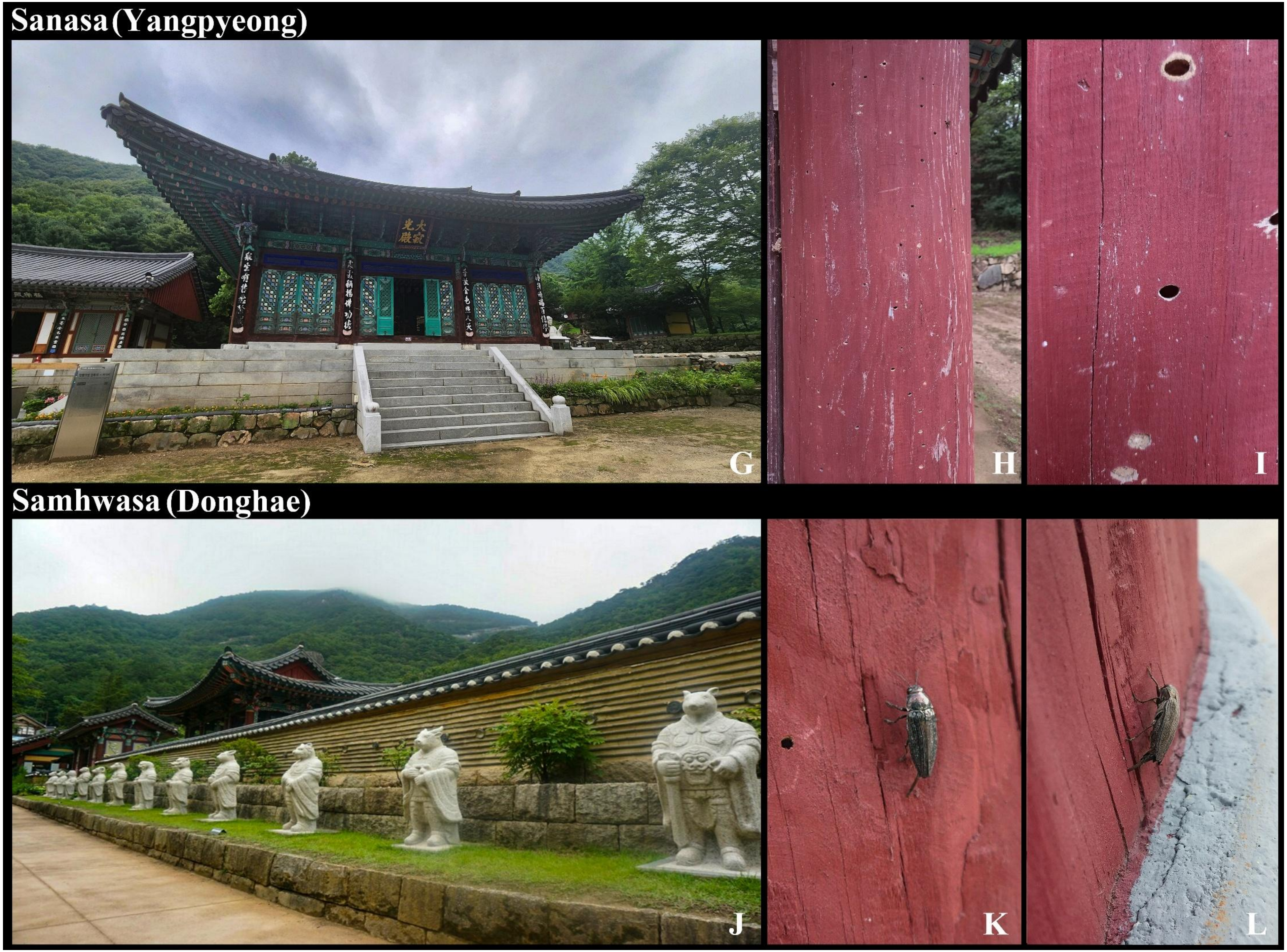
Sanasa (Yangpyeong)
G
H
I
Samhwasa (Donghae)
J
K
L

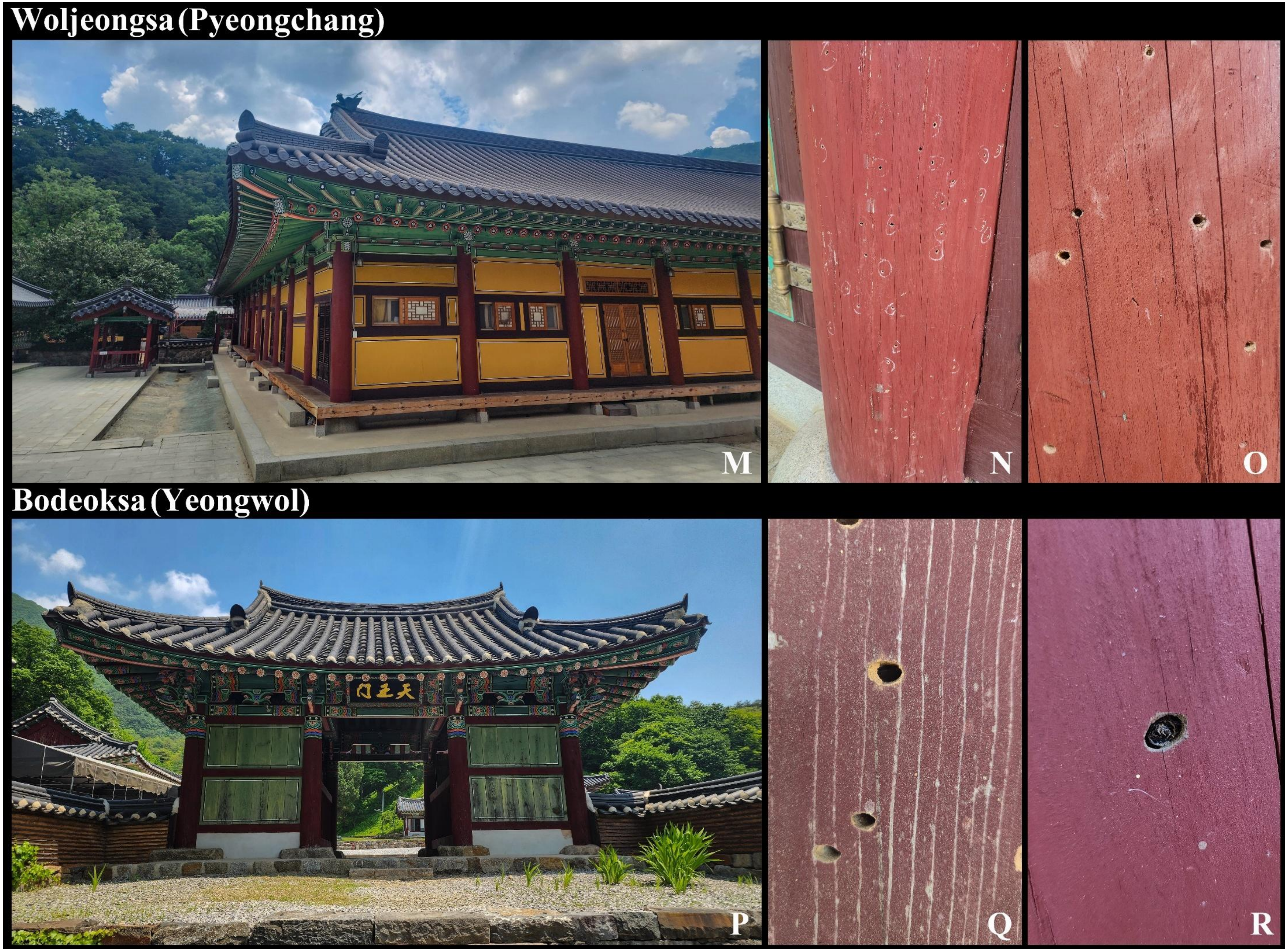
Woljeongsa (Pyeongchang)
M
N
O
Bodeoksa (Yeongwol)
P
Q
R

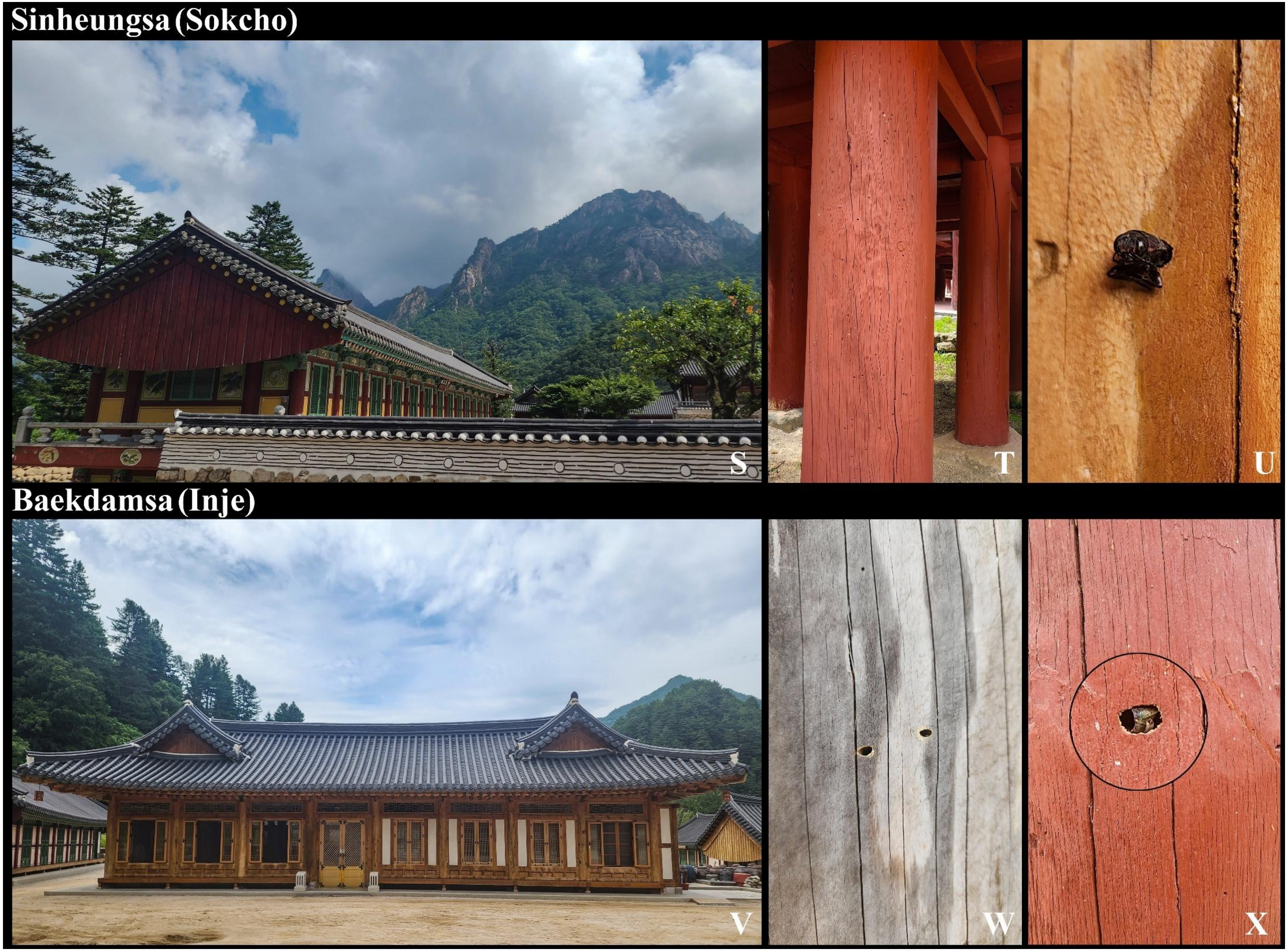
Sinheungsa (Sokcho)
S
T
U
Baekdamsa (Inje)
V
W
X

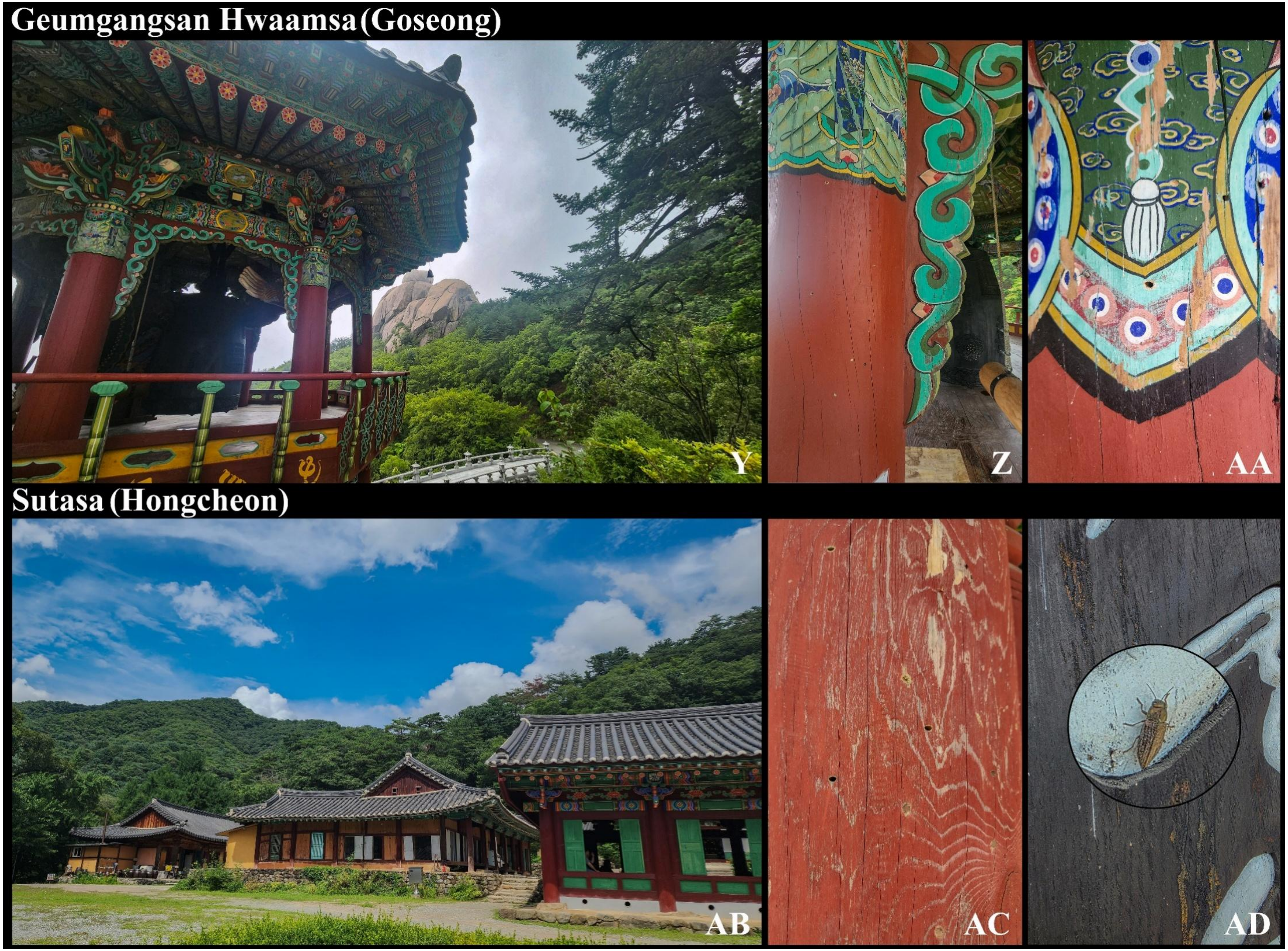
Geumgangsan Hwaamsa(Goseong)
Y
Z
AA
Sutasa (Hongcheon)
AB
AC
AD

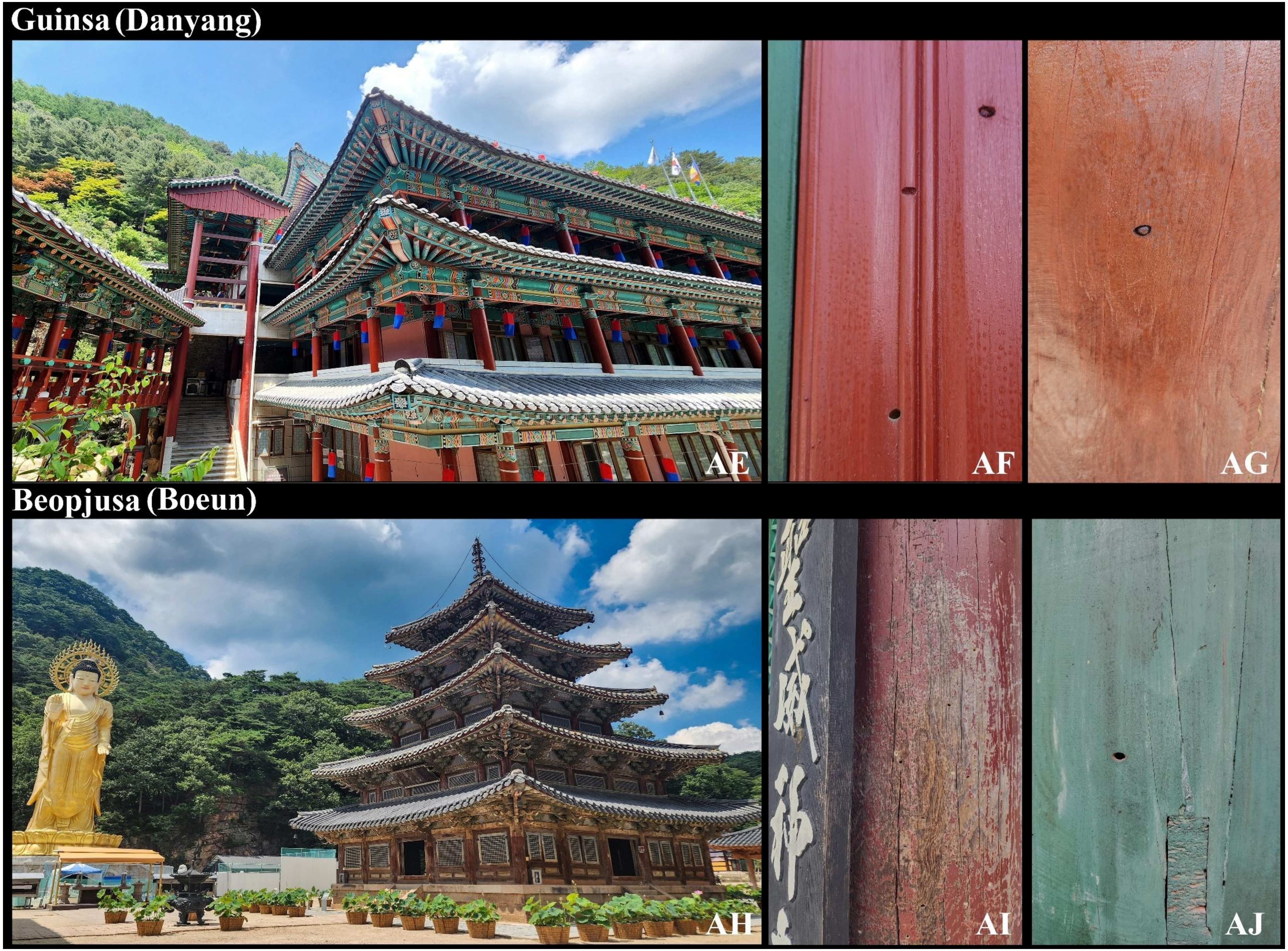
Guinsa (Danyang)
AE
AF
AG
Beopjusa (Boeun)
AH
AI
AJ

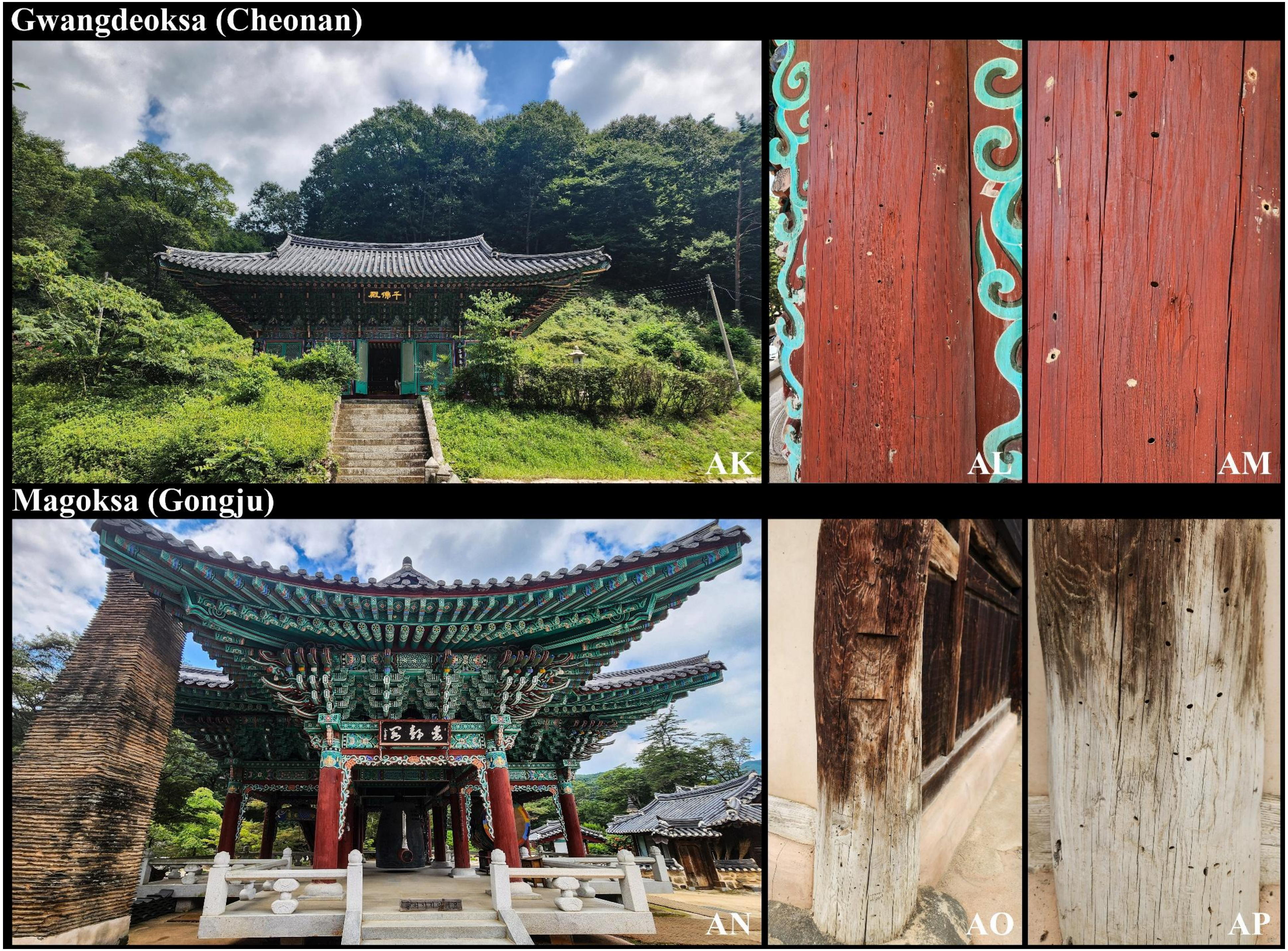
Gwangdeoksa (Cheonan)
AK
AL
AM
Magoksa (Gongju)
AN
AO
AP

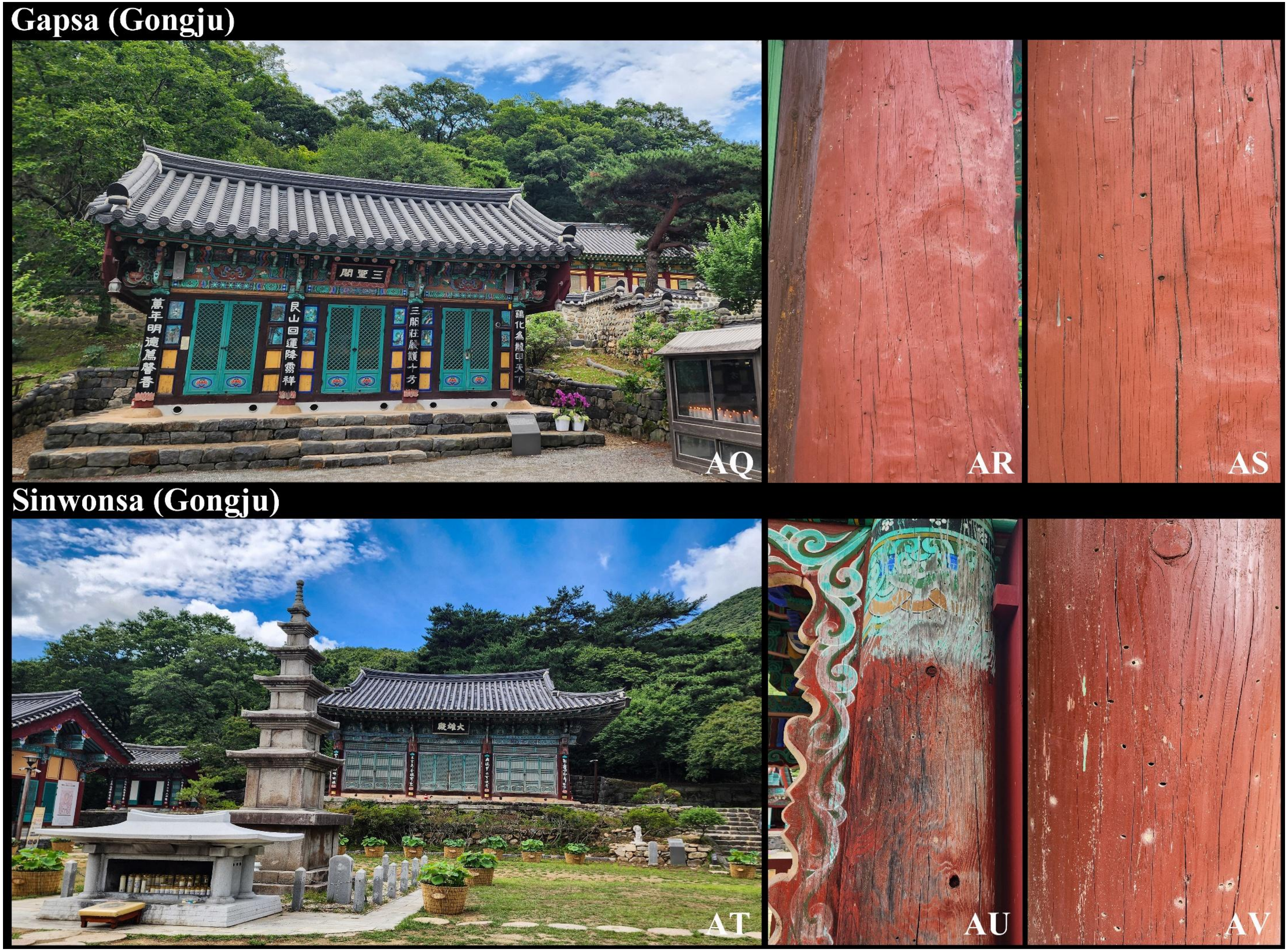
Gapsa (Gongju)
AQ
AR
AS
Sinwonsa (Gongju)
AT
AU
AV

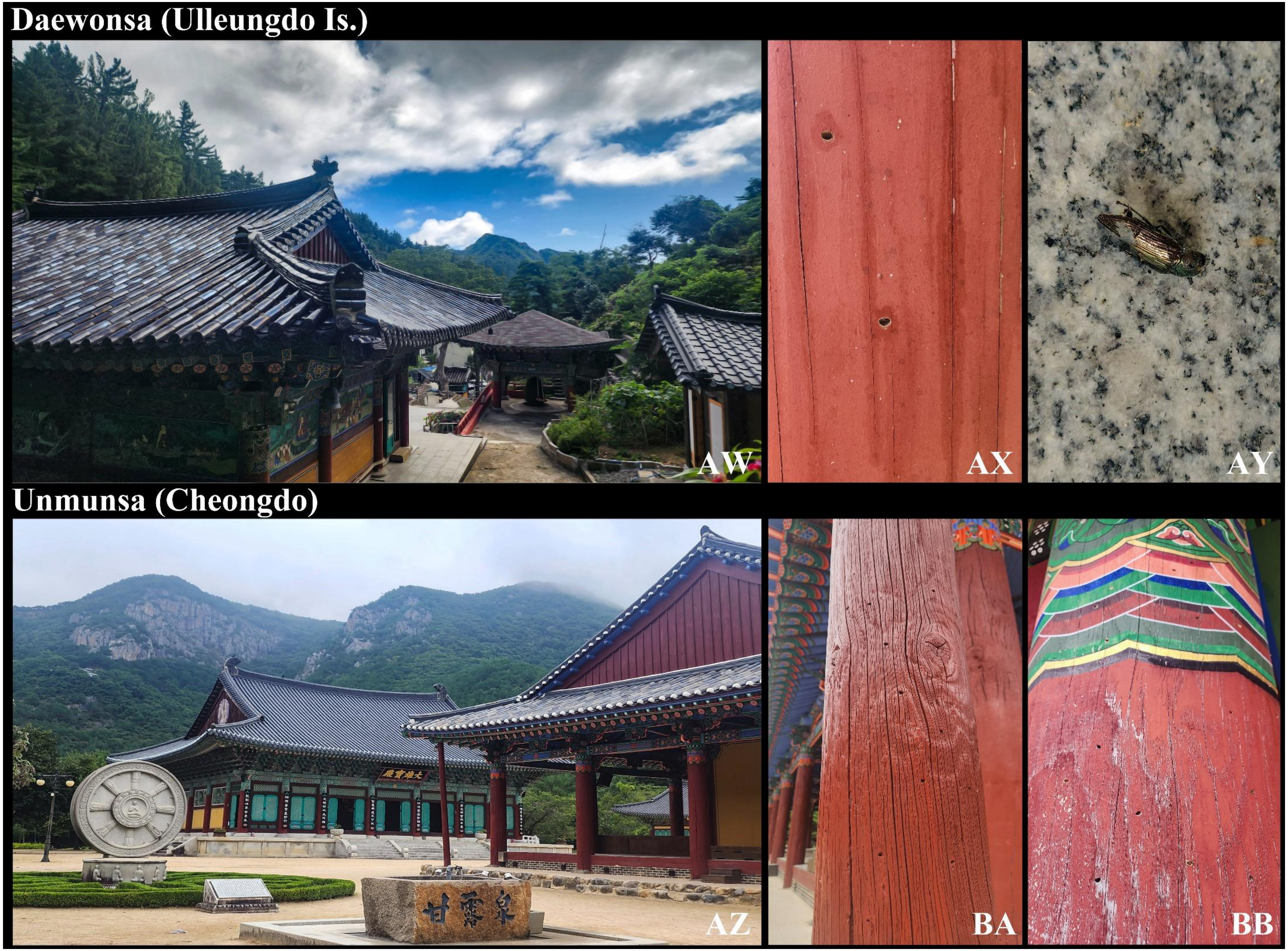
Daewonsa (Ulleungdo Is.)
AW
AX
AY
Unmunsa (Cheongdo)
甘露泉
AZ
BA
BB

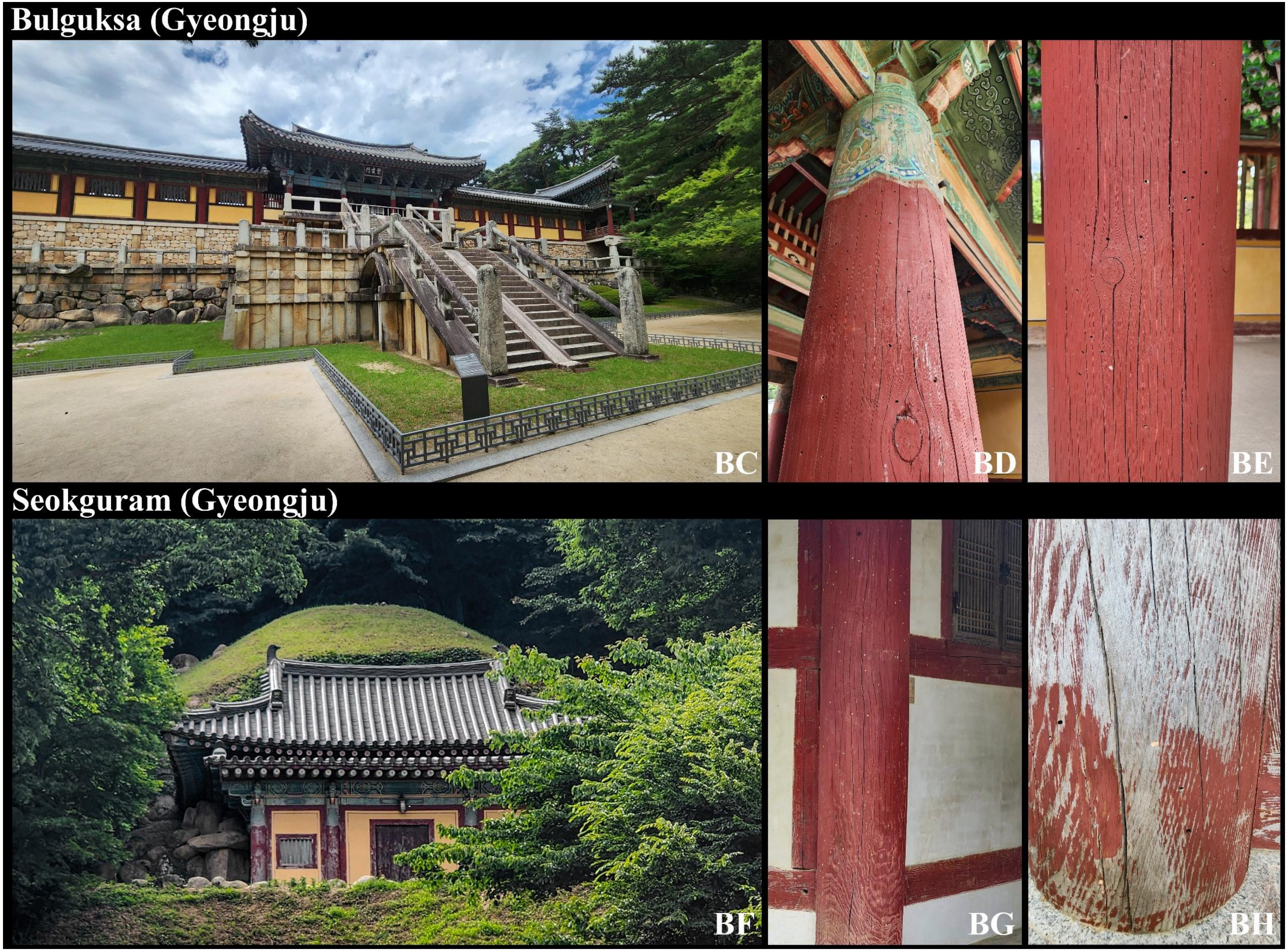
Bulguksa (Gyeongju)
BC
BD
BE
Seokguram (Gyeongju)
BF
BG
BH

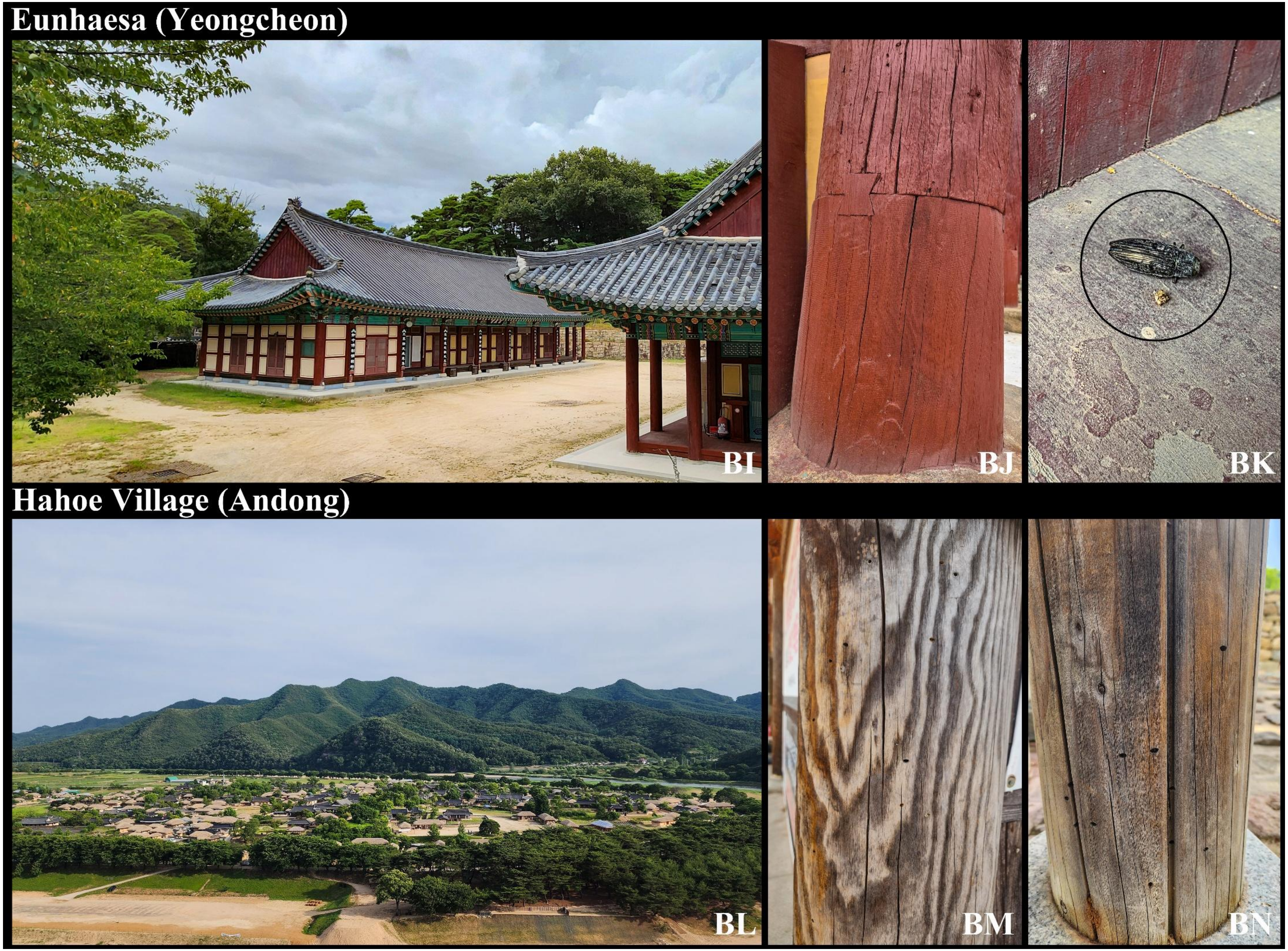
Eunhaesa (Yeongcheon)
BI
BJ
BK
Hahoe Village (Andong)
BL
BM
BN

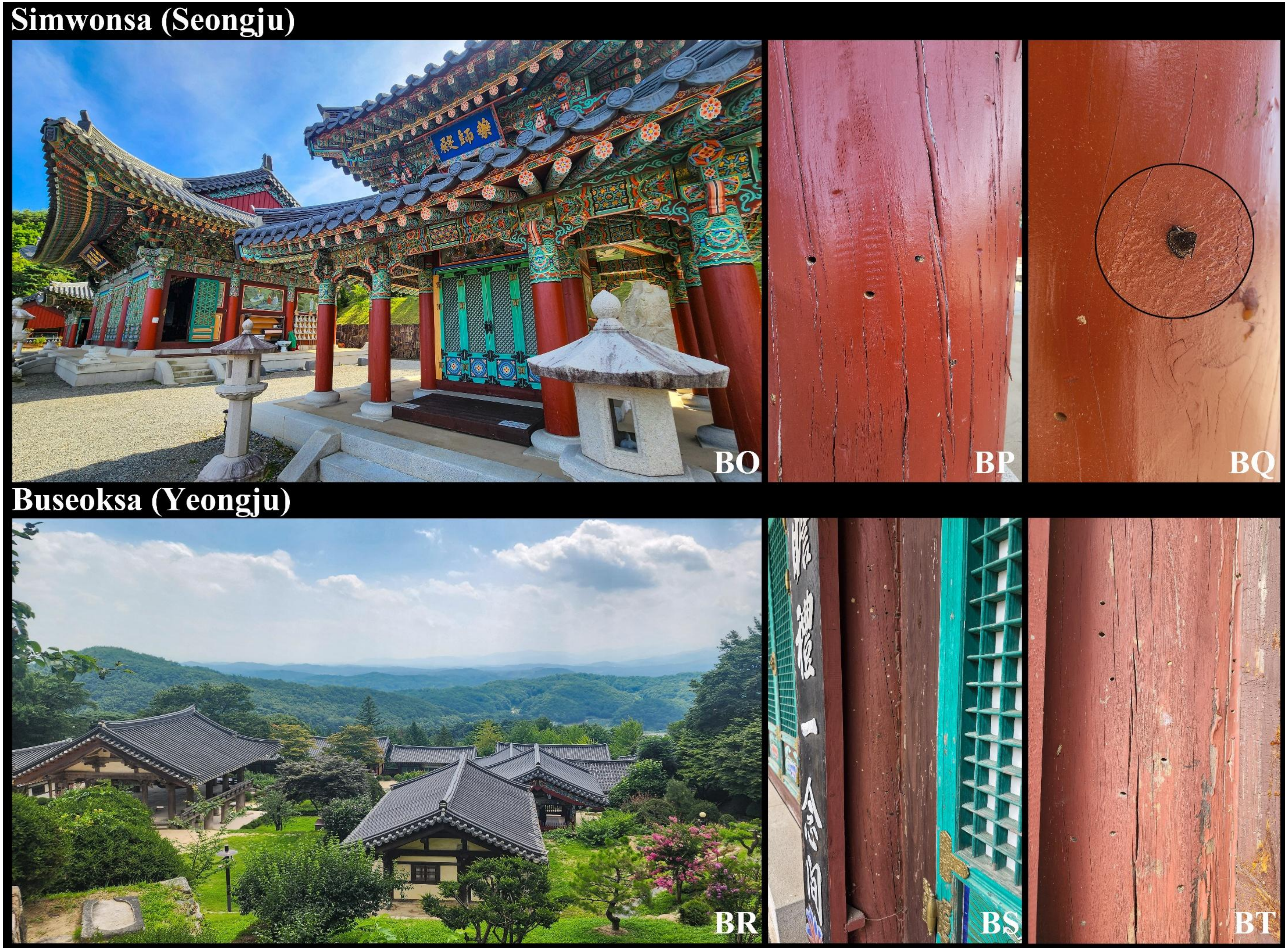
Simwonsa (Seongju)
BO
BP
BQ
Buseoksa (Yeongju)
BR
BS
BT

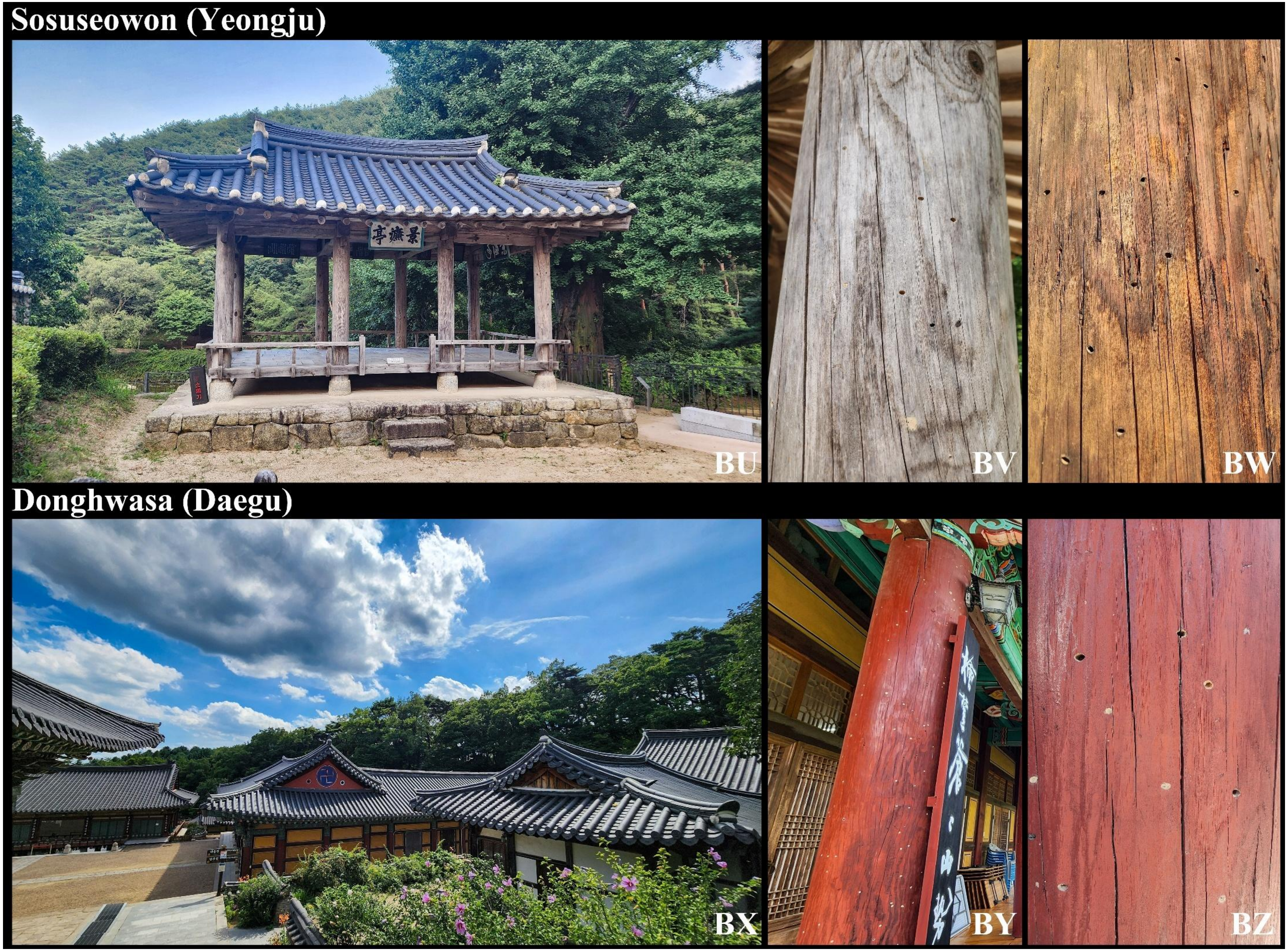
Sosuseowon (Yeongju)
BU
BV
BW
Donghwasa (Daegu)
BX
BY
BZ

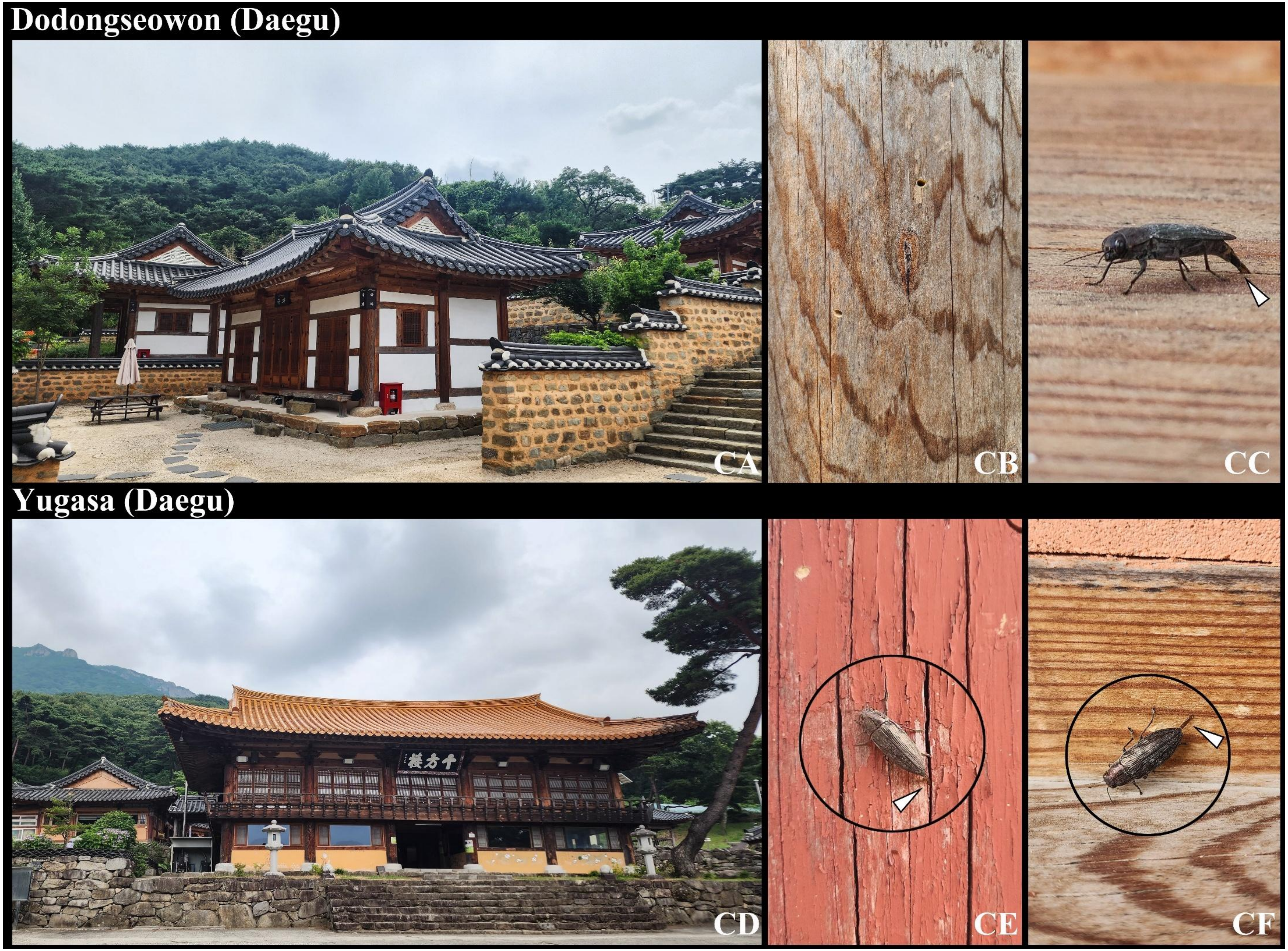
Dodongseowon (Daegu)
CA
CB
CC
Yugasa (Daegu)
CD
CE
CF

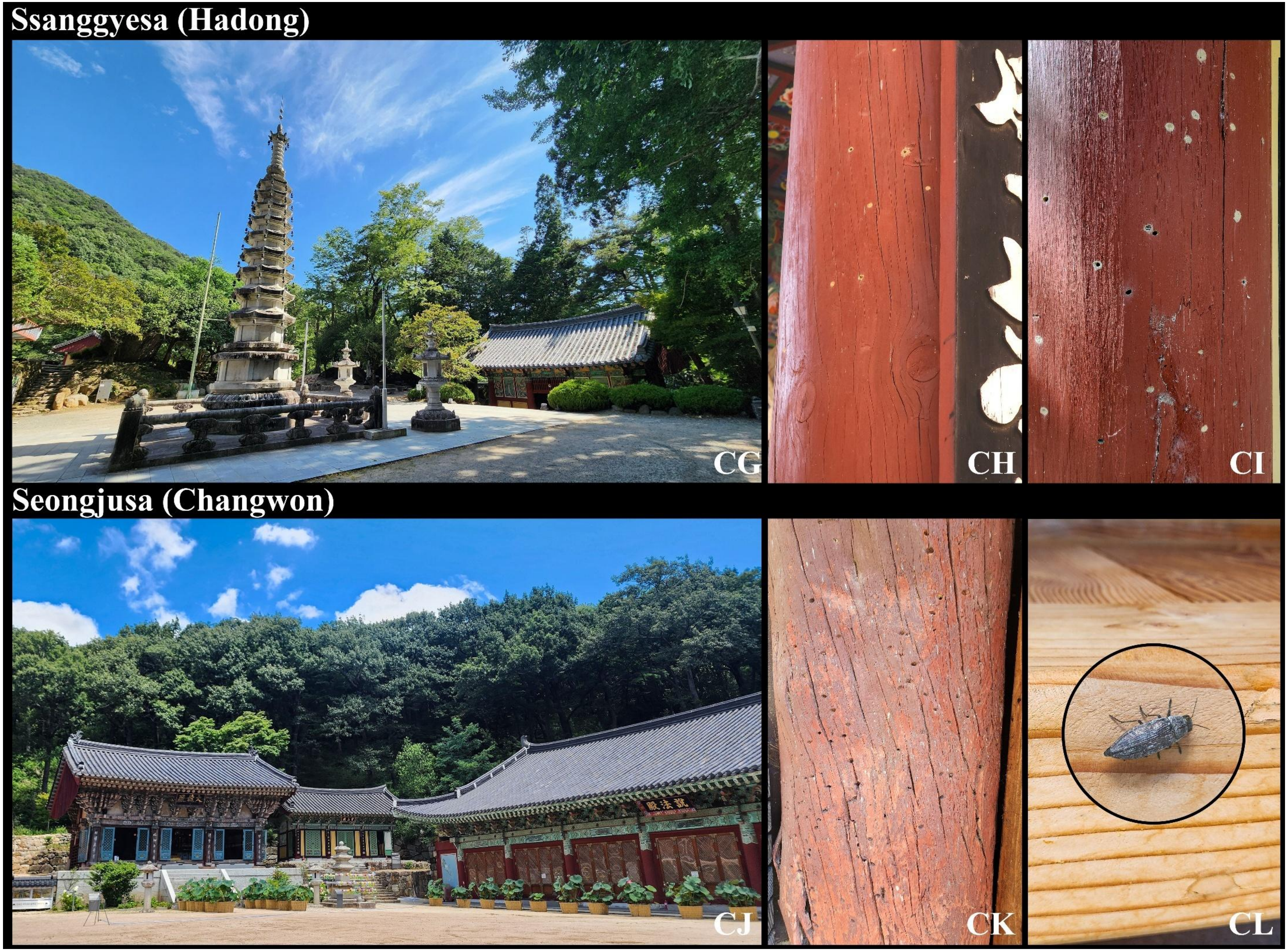
Ssanggyesa (Hadong)
CG
CH
CI
Seongjusa (Changwon)
CJ
CK
CL

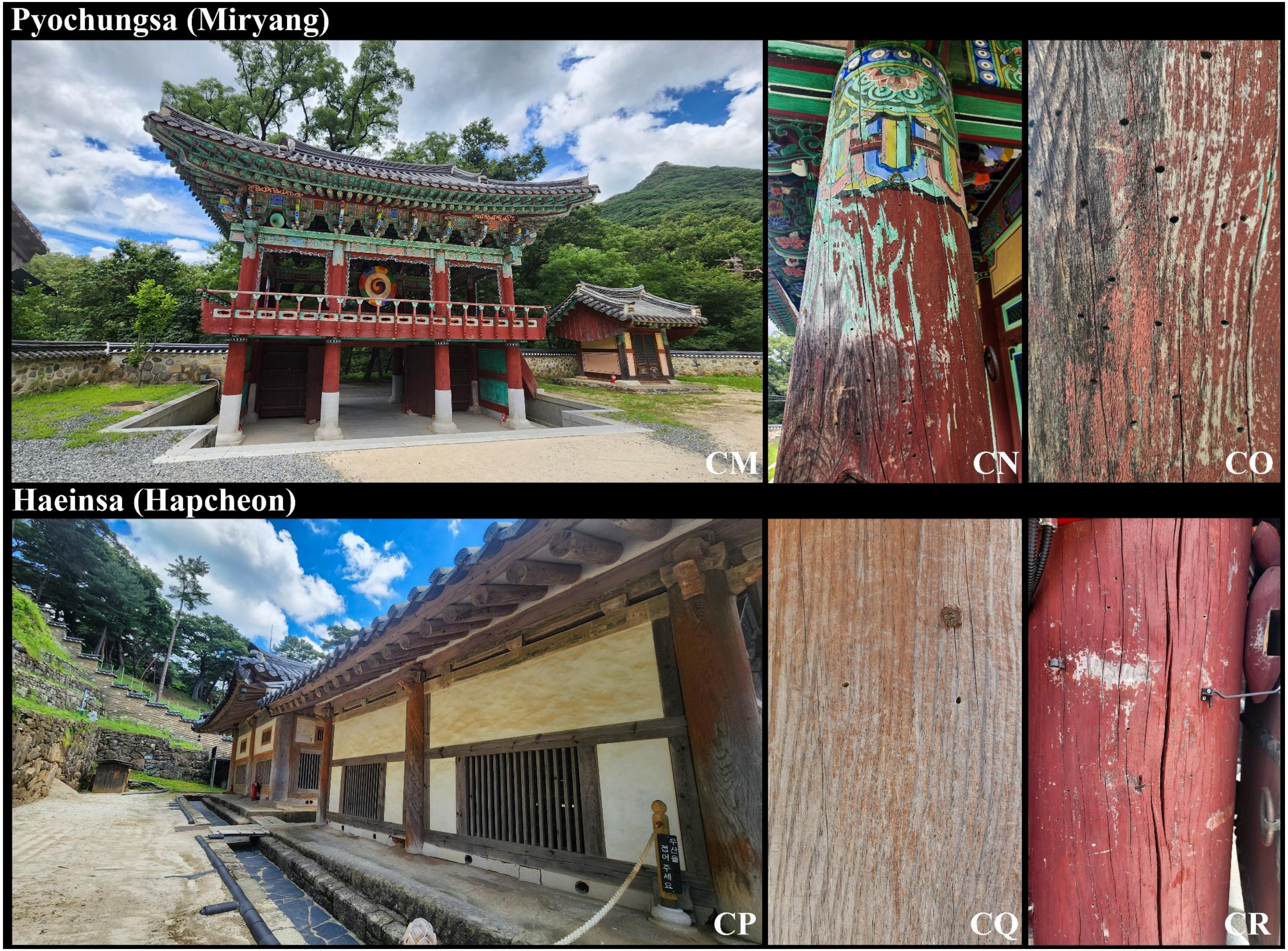
Pyochungsa (Miryang)
CM
CN
CO
Haeinsa (Hapcheon)
우산을
접어 주세요
CP
CQ
CR

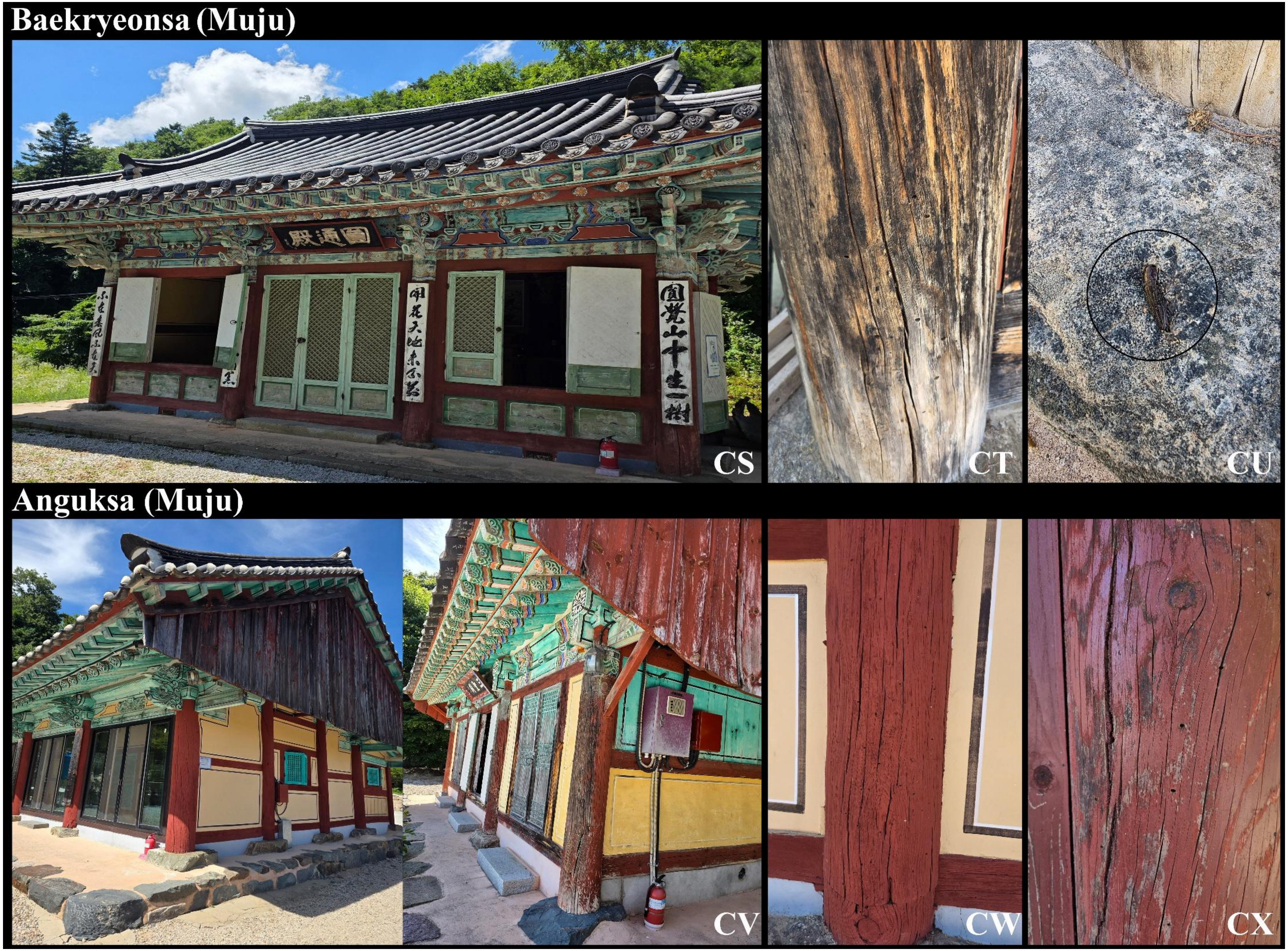
Baekryeonsa (Muju)
CS
CT
CU
Anguksa (Muju)
CV
CW
CX

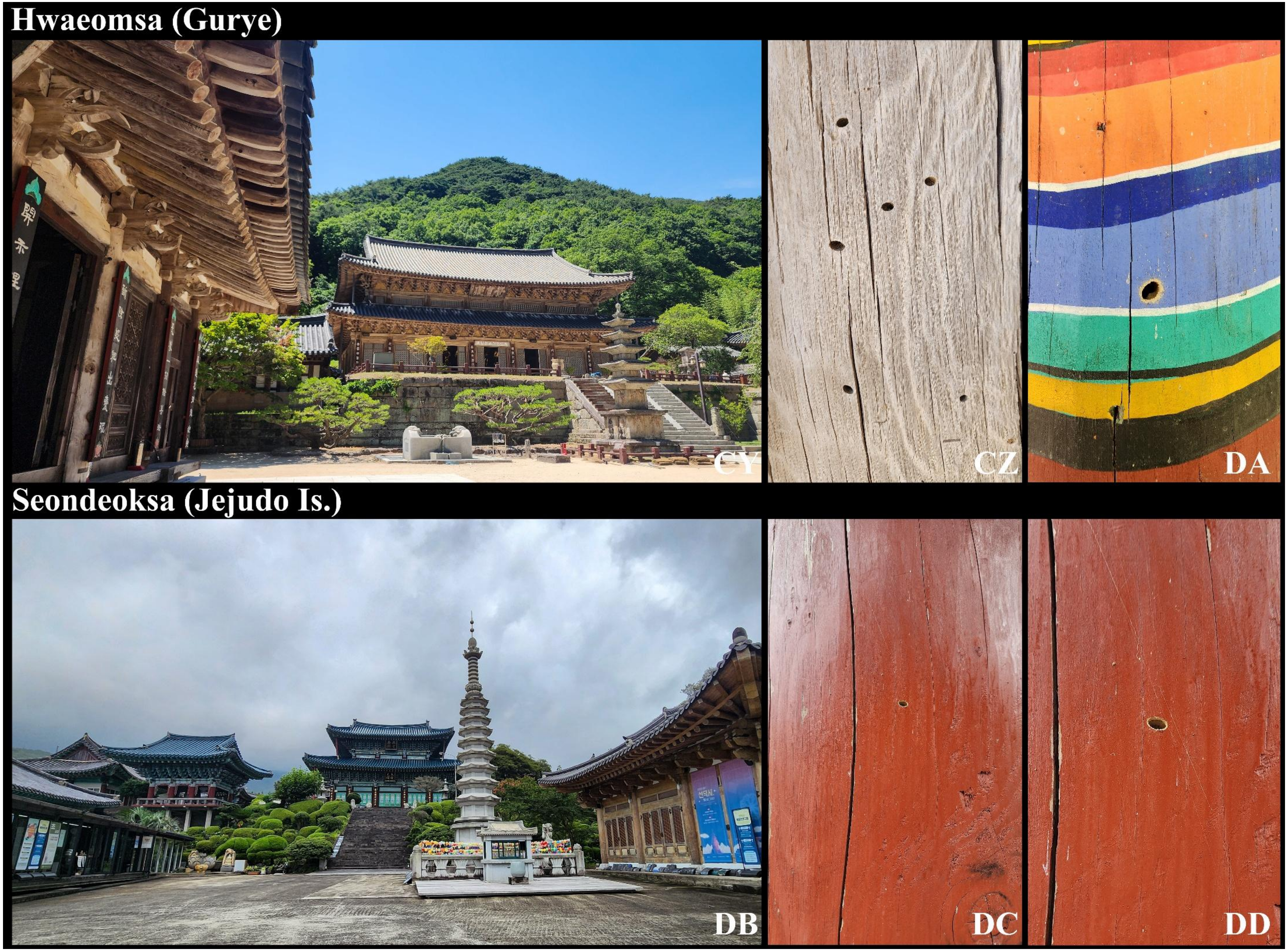
Hwaeomsa (Gurye)
CY
CZ
DA
Seondeoksa (Jejudo Is.)
DB
DC
DD
